# Structural basis of drug efflux by the staphylococcal efflux pump QacA

**DOI:** 10.64898/2026.04.10.717755

**Authors:** Léni Jodaitis, Patrick Sutton, Andrew Hutchin, Abolfazl Dashtbani-Roozbehani, Kyo Coppieters, Els Pardon, Jan Steyaert, Chloé Martens, Megan L. O’Mara, Melissa H Brown, Cedric Govaerts

**Affiliations:** Biochemistry and Structural Biology, Université Libre de Bruxelles, Brussels, Belgium; Australian Institute for Bioengineering and Nanotechnology, The University of Queensland, St Lucia 4072, Australia; College of Science and Engineering, Flinders University, Bedford Park, Australia; Structural Biology Brussels, Vrije Universiteit Brussel (VUB), Brussels, Belgium; VIB-VUB Center for Structural Biology, VIB, Brussels, Belgium, Brussels, Belgium; ARC Training Centre for Biofilm Research and Innovation, Flinders University, Bedford Park, Australia

## Abstract

The QacA DHA2 exporter from *Staphylococcus aureus* is a prototypical multidrug transporter that, like other bacterial efflux pumps, can extrude a wide range of cytotoxic compounds thus playing a crucial role in antimicrobial resistance. Here, we report crystal structures of wild-type QacA in three key conformational states: inward-open, outward-open and ethidium-bound, representing the first ligand-bound structure of a 14 transmembrane helices (TM) DHA2 transporter. In combination with computational and functional studies, these structures provide a mechanistic framework to understand drug recognition and extrusion. Structural analyses reveal remarkable adaptability within the binding pocket, including a ligand-induced deformation of TM5 that enables coordination of ethidium bromide in the outward-open state. Molecular dynamics simulations show spontaneous lipid entry into the transporter core and suggest that substrate binding from the inner membrane leaflet initiates a conformational transition to an outward-open state, stabilizing high-affinity interactions. Subsequent binding site protonation drives substrate extrusion. Together, these findings elucidate the structural dynamics and mechanistic underpinnings of QacA-mediated multidrug transport, highlighting conformational flexibility and proton-coupled electrostatic changes as key determinants of multidrug recognition and extrusion. This study provides a foundational framework for developing targeted inhibitors to combat bacterial multidrug resistance.

## INTRODUCTION

With current estimates of more than 1 million deaths per year directly attributable to and about 5 million associated with infections by resistant bacteria [1], antimicrobial resistance (AMR) represents a formidable public health challenge for the coming years. Drug efflux is a key mechanism in AMR as it secures bacterial survival following an initial chemical stress, thus enabling genetic adaptation that will provide adaptive resistance to the cytotoxic pressure. Expression of multidrug transporters, which extrude chemically diverse antibiotics, can be rapidly upregulated in pathogens under infection-related stresses (such as exposure to serum or blood) [2], underscoring importance of this key resistance mechanism in the clinical setting.

Multidrug exporters represent a scientific enigma as they depart from the classical conundrum that a given protein/enzyme/transporter has specifically evolved recognition of one or few substrates. Indeed, these transporters can not only recognize a large spectrum of chemicals, but also efficiently expel them outside the cells. This implies that the structural adaptation must go beyond versatility in binding and will involve adjustments of the energy-dependent conformational changes required for extrusion.

A remarkable example is QacA, a multidrug transporter expressed by *Staphylococcus aureus*, the deadliest Gram-positive pathogen on a global scale [3]. QacA can efficiently expel over 30 monovalent and bivalent cationic, lipophilic antimicrobials, including antiseptics and disinfectants like benzalkonium and chlorhexidine [4]. Due to its presence on plasmids, this efflux pump can swiftly disseminate among staphylococci in both clinical settings and natural environments [5]. QacA is a transporter of the Major Facilitator Superfamily (MFS) and belongs to the DHA2 (drug/H+ antiporter) sub-family. This sub-family is characterized by having 14 transmembrane helices (TM) in contrast to the 12-TM architecture typical of canonical MFS transporters. In DHA2 members, the two additional TM helices are located on the side of the core 12-helix bundle, forming an extended structure distinctive to this sub-family.

MFS transporters form an extraordinarily large family of secondary transporters including various types of importers and exporters and are found in all kingdoms of life, as recorded in the Transporter Classification Database TCDB [6]. They have been extensively studied and current transport models suggest that the pseudo-symmetrical organization of the transmembrane helical bundle enables a rocker-switch motion between the N-terminal and C-terminal lobes to allow for alternating access of either side of the membrane [7]. Specifically, the transporters are alternating between an *outward-open* (or outward-facing) state where the binding pocket is exposed to the extracellular space (or periplasmic space in the case of Gram-negative bacterial transporters) and an *inward-open* (or inward-facing) state where the binding pocket is exposed to the cytosol. Structural studies show that, overall, these states are related by a rigid body rotation of the N- and C-lobes and that each conformation is stabilized by sets of polar interactions connecting the two lobes either on the intracellular or extracellular end [7]. Transitioning from one state to the other would require overcoming such energetic barriers, for example through ligand binding.

Notably, dynamics studies have shown that the transition mechanism between inward- and outward-open states may not follow a strict, global rigid-body rotation but, rather, that local reorganization may be involved in the transition state(s) [8] thus suggesting that local flexibility may be required for the conformational switch.

For multidrug transporters, the versatility of substrates implies adaptation of the binding pocket to the structural diversity of substrates. In the case of another MFS drug pump, LmrP, the presence of a lipid molecule inside in the binding pocket was proposed to provide such conformational diversity along with appropriate hydrophobicity [9]. Similarly, in the MFS efflux pump EfpA from *Mycobacterium tuberculosis*, cryo-EM structures show multiple lipid molecules occupying a central cavity, suggesting a role in lipid translocation [10]. However, such embedded lipids have not been described in the structures of other major classes of drug exporters, including RND and ABC transporters, suggesting that different mechanisms may underlie substrate promiscuity across transporter families.

Here, we solved the structures of QacA in both outward-open and inward-open states, and have provided the first structure of a DHA2 transporter bound to a substrate. These structures reveal the key interactions regulating the conformational equilibrium but also identify unpredicted conformational plasticity in the binding pocket that enables recognition of the substrate.

## RESULTS

### Structure of the outward-open state

Wild-type (WT) QacA was expressed in *Escherichia coli* and purified to homogeneity (see Methods). To obtain crystals, the protein was incubated with exogenous lipids prior to crystallization trials. Diffracting crystals were obtained under different conditions (e.g. without substrate, bound to a nanobody or to EtBr), leading to high-resolution structure of QacA in different conformations. Crystals in the apo state diffracted up to 2.8 Å in the best direction (see Table S1) and the structure was solved by molecular replacement using the Alphafold prediction as a search model.

The refined structure of QacA shows the canonical MFS pseudo-symmetrical 12 TM bundle, containing the N- and C-lobe (each composed of 6 helices) along with the two extra TM helices found in the DHA2 sub-family of drug exporters as well as other 14 TM MFS transporters (Figure 1A, 1B). In addition, QacA bears a structured extracellular loop 7 (ECL7) between TM13 and 14 composed of a small 4-helix bundle. Additional densities decorate the protein where either lipids or detergent molecules can be modelled (Figure S1).

**Figure 1.**
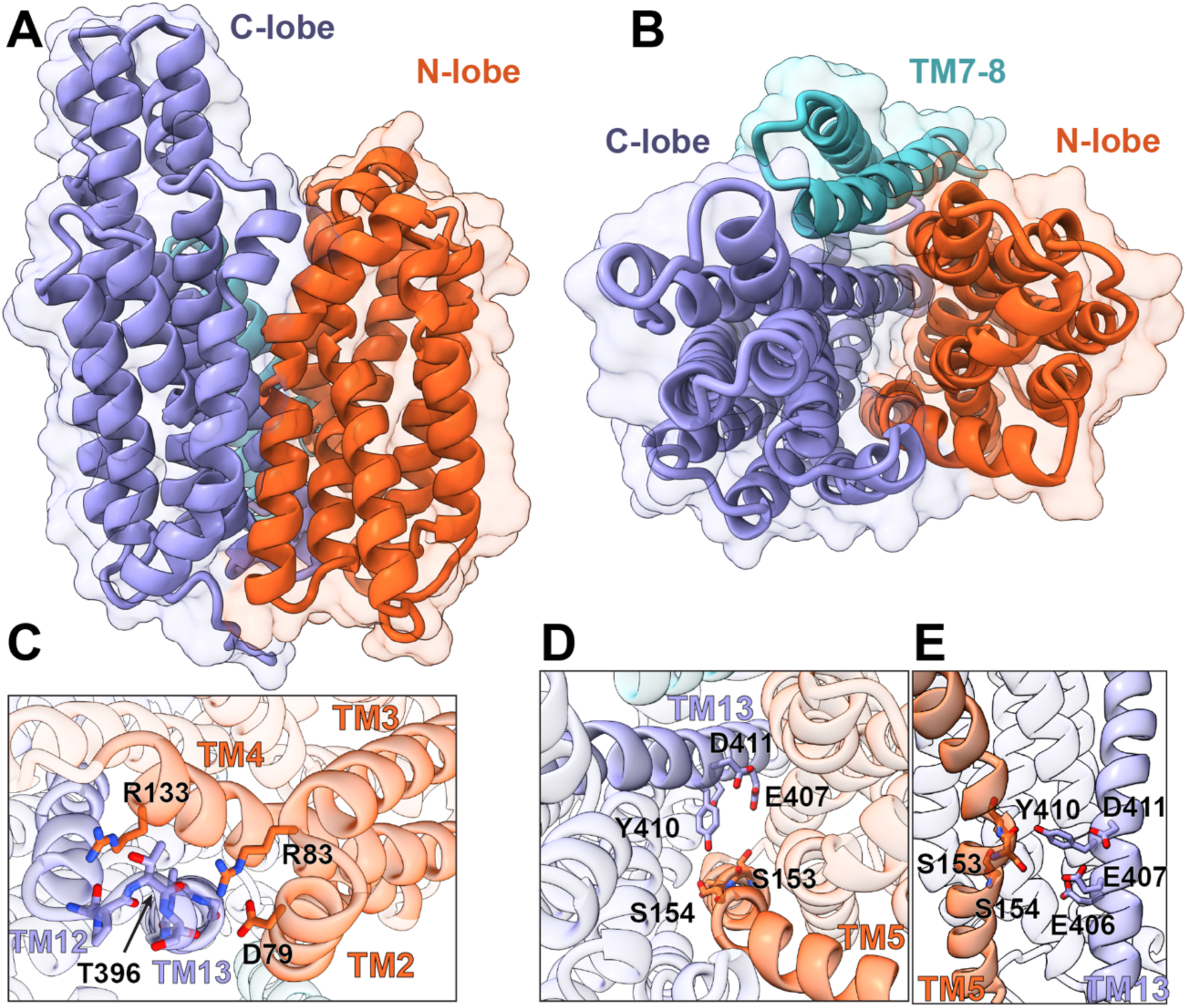
Crystal structure of QacA in outward-open conformation. A and B: View from the side and extracellular side of the backbone structure of QacA represented as ribbons. The first six helices (N-lobe) are colored in orange and the last six helices (C-lobe) are colored in purple while helices 6 and 7 (specific to 14-TM MFS transporters) are shown in cyan. The semitransparent surface gives the molecular volume of the protein. C: Close-up view of motif A, connecting the N- and C-lobe at the bottom of TM2, TM3, TM4, TM12 and TM13. D and E: Top and side view of the motif Y (see text) connecting the N- and C-lobe via Y410.

In this apo state, QacA adopts an outward-open conformation where the expected ligand binding pocket (i.e. the crevice between the N- and C-lobes) is clearly solvent-accessible to the extracellular side and is similar to the conformation of a functionally-inactive mutant of QacA recently solved by cryo-EM at 3.6 Å [11]. The crystallographic and cryo-EM structures differ by 1.2 Å root mean squared deviation (RMSD), with structural differences located primarily in the inter-helical loops (RMSD of 0.8 Å between the helical regions).

We assessed the crystal structure using molecular dynamics (MD) simulations in a lipid membrane (see Methods) which show that the conformation observed in the crystal is stable with an RMSD of 2.2 ± 0.3 Å calculated across the combined 1500 ns of simulation (Figure S2).

The crystal structure shows a variety of polar interaction networks that appear to stabilize the outward-open conformation. At the intracellular end, QacA bears the conserved so-called motif A (GX3-(D/E)-(R/K)-X-G-[X]-(R/K)-(R/K) [12]) which links intracellular loop 1 (ICL1) with the bottom of TM13 (Figure 1C) Structures of related MFS transporters, such as MdfA [13] or LmrP [9] show that the conserved aspartate connects the intracellular ends of TM2 on the N-lobe and TM11 on the C-lobe (TM13 in transporters with 14TMs such as QacA) by interacting with the backbone nitrogen atoms at the bottom of TM11 thus stabilizing the inward-closed/outward-open state. Protonation of this conserved aspartate has been proposed to act as a switch enabling the conformational changes to the inward-open state [14]. In QacA, this connecting motif is partially modified (the starting glycine residues are not present) and while the aspartate (D79) is pointing towards TM13, it shows direct interactions between R83 (TM3) of motif A and the carbonyl of T396, as well as between the sidechain of R133 (TM4) and the backbone carbonyl of T394 (TM14). This indicates that, although partially conserved, motif A has evolved in QacA, with potential structural and functional consequences. Mutagenesis of either R83 or R133 to cysteine leads to reduced activity toward ethidium and the bivalent substrates, supporting the functional role of these residues (Table 1). In MD simulations of the outward-open state, an extended electrostatic network is present across the charged residues of Motif A for over 97 % of the 1500 ns of simulations. Here the guanidino side chain of R133 interacts with the backbone carbonyl and sidechain hydroxyl of T394 for the entire 1500 ns of simulations, while the R83 guanidino side chain forms a salt bridge with the carboxylic sidechain of D79, which also interacts with the backbone amine of G400 (Figure S3).

**Table 1.** Antimicrobial resistance profiles of QacA mutants.

| Mutation | Location <sup>a</sup> | Expression<br>(% of WT) | MICs of QacA mutants (% WT)<sup>b</sup> |  |  |  |  |  |
| --- | --- | --- | --- | --- | --- | --- | --- | --- |
|  |  |  | Monovalent substrates |  |  | Bivalent substrates |  |  |
|  |  |  | Et<sup>c</sup> | R6G | BK | DQ | CH | PE |
| WT QacA<sup>d</sup> |  | 100 | 100 (250) | 100 (800) | 100 (100) | 100 (300) | 100 (7) | 100 (240) |
| No QacA |  | NA | 10 | 31 | 30 | 25 | 14 | 33 |
| V31C | TM1 | 100 | 68 | 49 | 107 | 88 | 54 | 88 |
| D34C | TM1 | 90 | 10 | 45 | 100 | 25 | 8 | 50 |
| M35C | TM1 | 94 | 20 | 45 | 75 | 50 | 50 | 50 |
| P43C | TM1 | 100 | 80 | 63 | 75 | 75 | 75 | 75 |
| D61C | TM2 | 50 | 110 | 80 | 114 | 85 | 79 | 75 |
| D79C | TM2 | 84 | 140 | 125 | 100 | 80 | 100 | 100 |
| R83C | Loop 2-3 | 130 | 55 | 90 | 89 | 45 | 75 | 60 |
| R133C | Loop 4-5 | 100 | 83 | 100 | 111 | 100 | 100 | 100 |
| K140A | Loop 4-5 | 105 | 40 | 47 | 50 | 40 | 27 | 41 |
| W149C | TM5 | 84 | 40 | 83 | 20 | 80 | 85 | 53 |
| S153C | TM5 | 125 | 46 | 94 | 110 | 73 | 85 | 60 |
| S153A | TM5 | 108 | 50 | 89 | 100 | 75 | 80 | 73 |
| S154C | TM5 | 85 | 70 | 89 | 90 | 80 | 70 | 83 |
| S154A | TM5 | 90 | 90 | 107 | 110 | 92 | 100 | 107 |
| K228A | Loop 7-8 | 80 | 80 | 74 | 67 | 67 | 82 | 65 |
| S231A | Loop 7-8 | 130 | 120 | 105 | 117 | 113 | 109 | 100 |
| K256A | TM8 | 87 | 120 | 110 | 117 | 107 | 127 | 100 |
| K272A | TM9 | 75 | 40 | 47 | 33 | 40 | 27 | 41 |
| M287C | TM9 | 95 | 50 | 46 | 80 | 54 | 100 | 71 |
| M287A | TM9 | 105 | 30 | 38 | 80 | 45 | 100 | 50 |
| Q302C | Loop 9-10 | 100 | 90 | 85 | 100 | 109 | 100 | 100 |
| Q302A | Loop 9-10 | 100 | 80 | 85 | 80 | 100 | 100 | 100 |
| M319C<sup>e</sup> | TM10 | 114 | 60 | 80 | 100 | 100 | 100 | 64 |
| M319A | TM10 | 110 | 80 | 58 | 80 | 91 | 100 | 93 |
| D323C | TM10 | 89 | 90 | 88 | 80 | 83 | 82 | 50 |
| R336A | Loop 10-11 | 88 | 120 | 105 | 108 | 93 | 100 | 76 |
| M380C<sup>e</sup> | TM12 | 94 | 100 | 100 | 100 | 100 | 71 | 83 |
| M380A | TM12 | 85 | 80 | 89 | 80 | 73 | 100 | 86 |
| S387C<sup>e</sup> | TM12 | 120 | 70 | 81 | 50 | 42 | 43 | 50 |
| E406C | TM13 | 105 | 20 | 46 | 90 | 50 | 80 | 67 |
| E407C | TM13 | 81 | 10 | 41 | 40 | 17 | 18 | 43 |
| Y410C | TM13 | 126 | 10 | 47 | 40 | 17 | 43 | 43 |
| D411C | TM13 | 118 | 10 | 47 | 40 | 100 | 100 | 36 |
| V415C | TMS13 | 110 | 128 | 77 | 226 | 96 | ND | 103 |
| S423A | TMS13 | 105 | 40 | 74 | 67 | 47 | 27 | 65 |

<sup>a</sup>Location of amino acid changes in QacA is based on the QacA structure (Figure 1).
<sup>b</sup>Numbers represent the relative resistance levels of QacA mutants compared to WT QacA, which is indicative of 100% resistance activity. MIC values were determined by a standard agar dilution method, where *E. coli* DH5α cells expressing WT and mutant QacA proteins were grown on LB agar supplemented with varying concentrations of each compound. All MIC data are representative of at least three independent experiments and are presented as percentage value relative to WT QacA. MIC values in **bold** indicate ≤50% of WT resistance, marking a functionally substantial reduction in QacA activity.
<sup>c</sup>Et: ethidium; R6G: rhodamine 6G; BK: benzalkonium; DQ: dequalinium; CH: chlorhexidine; PE: pentamidine.
<sup>d</sup>MIC value (μg mL<sup>-1</sup>) for WT QacA against each substrate is shown in parentheses.
<sup>e</sup>Data from previously published QacA mutants are included for comparison [19, 36].

A second polar network connects the N- and C-lobes at the level of the central binding pocket (Figure 1D, 1E). This network involves a motif located at the middle of TM13 composed of three negatively charged residues (E406, E407 and D411) and Y410, that we call here *motif Y*, and is conserved within a subclade of DHA2 proteins that includes QacA and several orthologs (Figure S4). Having three negatively-charged residues in close proximity within a TM bundle suggests a functional role. To support this, bacteria expressing QacA mutant derivatives of these residues, E406C, E407C, Y410C and D411C, exhibited substantially reduced resistance levels against most tested substrates compared to WT QacA (Table 1). Furthermore, the side chains of E407 and D411 are within 3.5 Å to each other and while a water molecule is observed nearby, protonation of one of these negatively-charged residues may be required for stability. Very clearly, Y410 is pointing directly towards the center of TM5, one turn below the conserved motif C (also known as the antiporter motif), which induces a strong kink break at the top of the helix [15, 16]. The reorientation of TM5 relative to TM10 during the transition to the inward-open state has been previously described [11]. However, in addition to this known conformational shift, our findings also reveal a structural connection between TM5 and TM13. Although Y410 does not appear to interact with specific side chains in TM5, it points towards the backbone carbonyl of S153 where, remarkably, the hydrogen bonding pattern of the helix is disrupted by a local bulge and thus available for interaction with the tyrosine sidechain. Notably, this local deformation in TM5 is not predicted by Alphafold. The crystal structure reveals possible electron densities for ions or water molecules suggesting the presence of a polar network bridging TM5 and TM13, but the limited resolution prevents definitive identification. Importantly, Y410 has been demonstrated as an essential residue of QacA, in line with the proposed structural (and functional) role suggested here [17].

Interestingly, in MD simulations, Na^+^ ions rapidly diffuse from the extracellular solution into the binding pocket and occupy the area adjacent to the TM5 bulge. Specifically, two Na^+^ ions bind to discrete binding sites (Na1 and Na2), where they are stably coordinated in an octahedral geometry (Figure 2). In each case, this involved polar bidentate coordination of Na^+^ mediated by protein hydroxyl or carbonyl groups and water. At the Na1 site, Na^+^ is coordinated by the hydroxyl groups of S150 and S154 (TM5), and three water molecules (Figure 2C). Here, water molecules take part in an interaction network with the carboxylate side chain of D323 (TM10), orienting it toward the center of the cavity. At the Na2 site, the second Na^+^ ion is stably coordinated by the carboxylate side chain of D34 (TM1), the Y63 (TM2) hydroxyl, the backbone carbonyl of V31 (TM1), and three water molecules (Figure 2D). Na^+^ ions occupied the Na1 (S154) and Na2 (D34) site for 81% and 86 % of the combined 1500 ns of simulation, respectively. Experimentally, D34 is important for the transport of some substrates examined, while D323 is critical for the transport of bivalent substrates [18] (Table 1).

**Figure 2.**
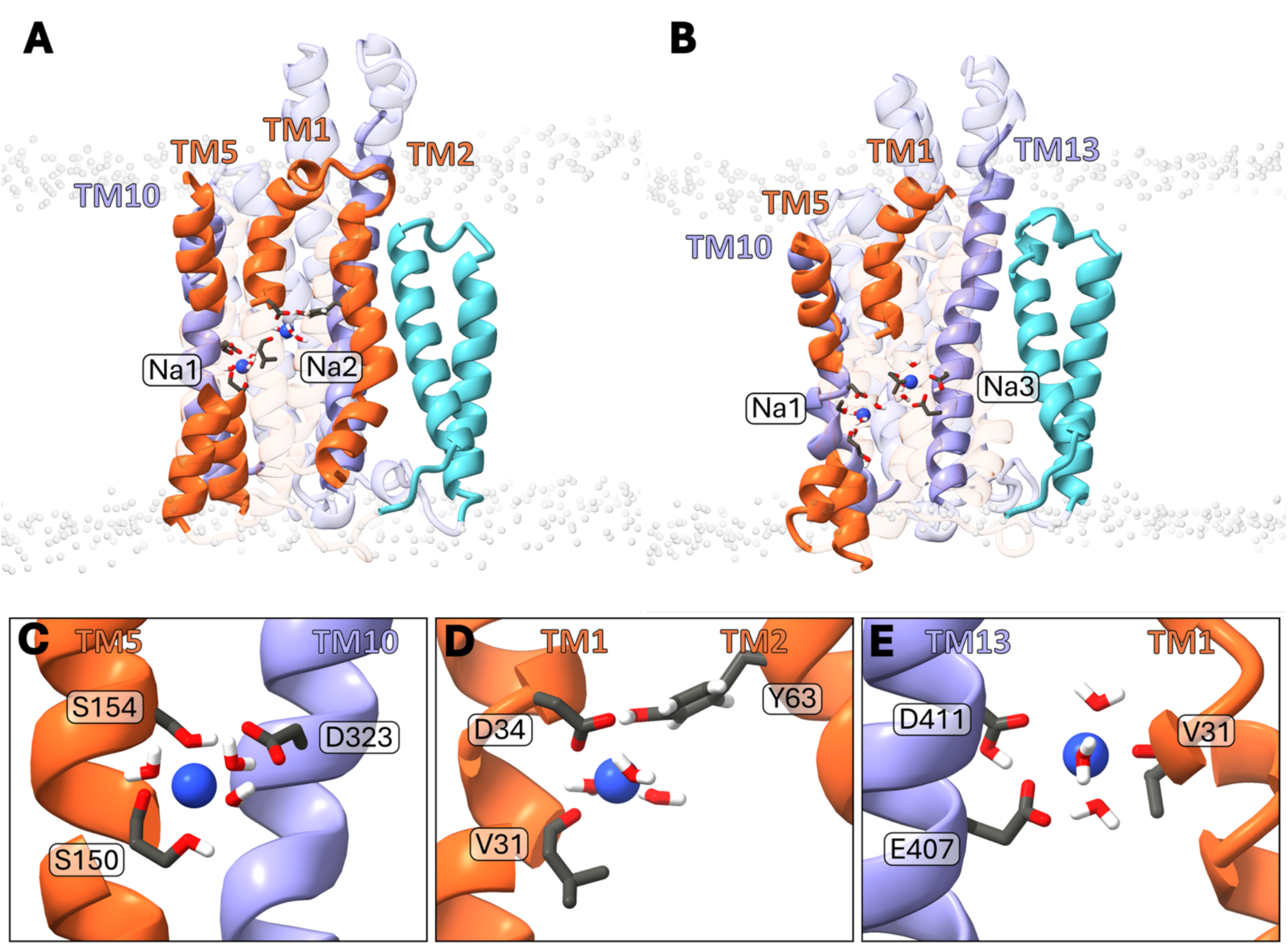
Sodium binding sites of inward and outward-open QacA in Molecular Dynamics simulations. Sodium binding sites of A: outward-open and B: inward-open QacA. C: The Na1 site is formed of S150, S154, and three water molecules, two of which are stabilized by D323. The Na1 site is present in both conformations of QacA. D: D34, V31, Y64 and three water molecules formed the Na2 site, which is only present in outward open QacA. E: Na3 is located adjacent to Na2, formed by D411, E407, V31 and three water molecules, present only in the inward-open conformation. Sodium ions are represented in dark blue.

### Inward-open nanobody-stabilized conformation of QacA

To stabilize different biologically relevant conformations of the transporter we generated QacA-specific nanobodies (see Methods) and isolated Nb89, a nanobody with nanomolar affinity against the protein (Figure S5). Crystals of QacA:Nb89 complexes were obtained which diffracted up to 3.3 Å. The structure was solved using an Alphafold-predicted template containing the sequences of both QacA and Nb89. In contrast to the apo structure, this nanobody-bound conformation is clearly in the inward-open state (Figure 3A). The nanobody makes several contacts with the structured ECL7 with an interface covering approximately 640 Å^2^ where both polar and hydrophobic interactions are found (Figure 3B). Compared to the outward-open conformation, the inward-open state requires a closing of the extracellular side of the TM bundle leading to direct interactions between ECL1 and ECL7 (which are separated by over 15 Å in the outward-open state). Binding of the nanobody appears to stabilize this ECL1-ECL7 interface as S59 and Y61 of Nb89 directly interact with ECL1, specifically with the sidechain of E50 and the backbone carbonyl of R47 (Figure 3C) while Y35 (of Nb89) coordinates an ionic bridge between R47 (ECL1) and E460 (ECL7) (Figure 3D).

**Figure 3.**
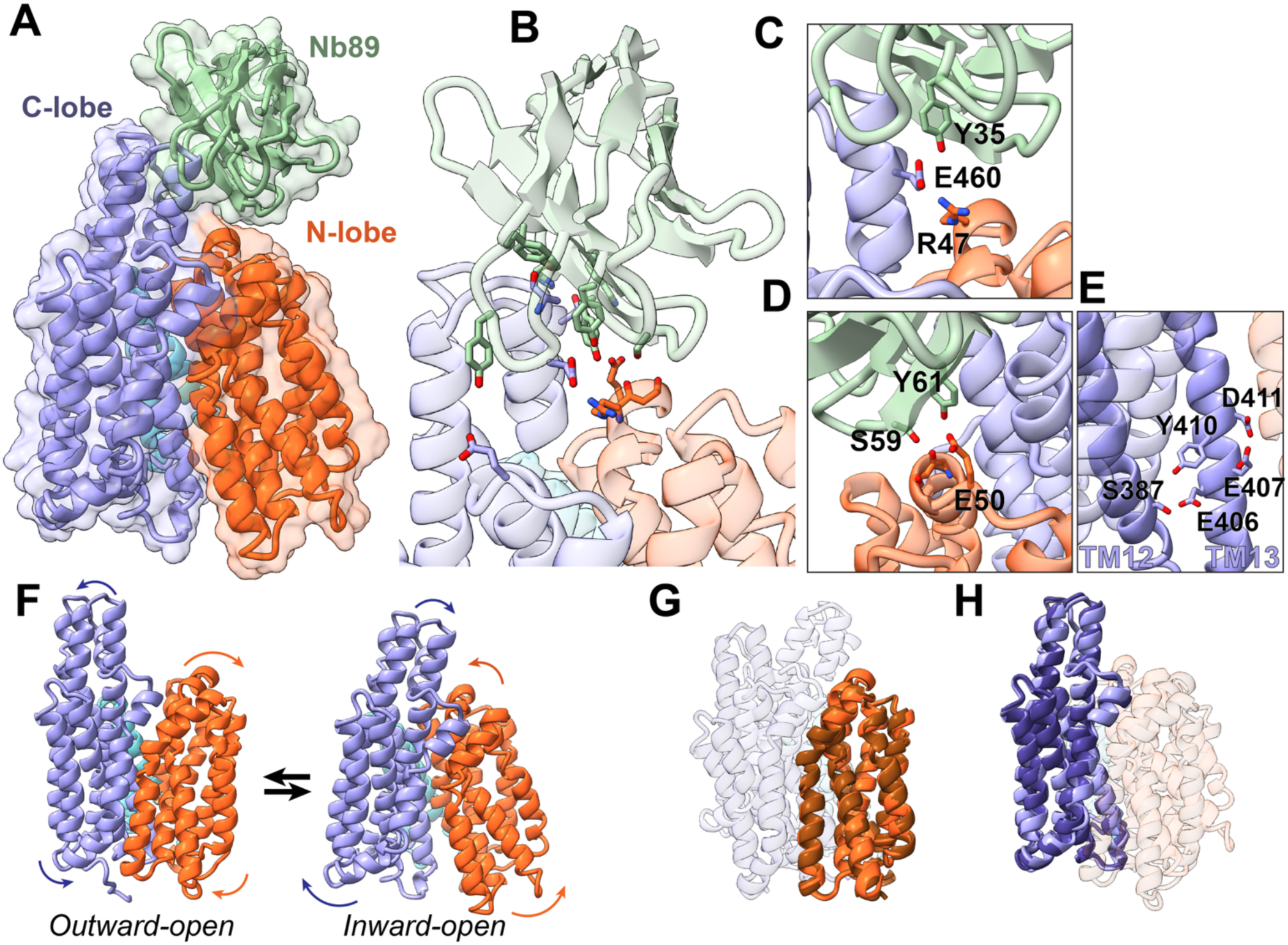
Crystal structure of nanobody stabilized inward-open QacA. A: View from the side and extracellular side of the backbone structure of QacA and Nb89 represented as ribbons. For QacA the first six helices (N-lobe) are colored in orange, the last six helices (C-lobe) are colored in purple and helices 6 and 7 (specific to 14-TM MFS transporters) are shown in cyan while the nanobody is in green. The semitransparent surface gives the molecular volume of the protein. B: Close-up view of interaction between the nanobody and QacA. C and D: detailed views of the polar interactions that appears to be involved in stabilizing the outward-closed conformation. E: Close-up view of the central Y motif showing the reorientation of Y410 withing the C-lobe. F: Ribbon representation showing side by side the outward-open and inward-open conformations (the nanobody is omitted for clarity). G and H: Superimposition of the two conformations on the N-lobe and C-lobe respectively, showing that the local structure of each lobe is similar in each state, which are thus related by a rigid-body motion.

Interestingly, motif Y, which links the N- and C-lobe in the outward-open state has rearranged in the inward-open conformation, with the side chain of Y410 pointing down and E406 making a polar interaction with S387 in TM12 (Figure 3E). This supports our previous studies which identified S387 as crucial for resistance to all bivalent substrates and the monovalent substrate benzalkonium [19].

Comparison of the two structures reveals that transition from outward-open to the inward-open states involves the closure of the extracellular side by about 10 Å while the intracellular side opens by about 15 Å (Figure 3F). As a consequence, solvent access to the central cavity is completely prevented from the extracellular side but, in turn, is readily possible from the intracellular all the way to the ligand binding pocket (as identified for ethidium, see below), including E406, E407 and Y410. This unilateral solvent accessibility is observed in both the crystal structure and throughout MD simulations of inward-open QacA with hydrophobic residues I39 and M40 (TM1), V60 (TM2), A291, L294 and L295 (TM9), M319 (TM10) and V418 (TM13), preventing water penetration to the extracellular space (Figure S3E). Among these, only mutation of M319 produces a moderate reduction in antimicrobial resistance (Table 1), while substitutions at the other residues had minimal impact and retained WT-like activity. These findings support the idea that M319 plays a particularly critical role in hydrophobic packing necessary to maintain the closed state of the extracellular side in the inward-open conformation.

The large-scale rearrangement relating the two states appears to solely involve a rigid-body motion of the N- and C-lobes relative to each other. The rocker-switch motion that relates the two conformations rotates around an axis located close to the center of the TM bundle (and thus the membrane). This leads to ample opening and closing on either side of the transporter thanks to strong helical kinks found in TM2, TM5, and TM10 which involve highly conserved proline residues. Indeed, the crystal structures reveal N-lobes from the inward-open and outward-open states are almost identical to each other (RMSD < 0.9 Å, Figure 3G), as are both C-lobes (RMSD <1 Å, Figure 3H). This is further supported by MD simulations of inward-open QacA. RMSDs were calculated for the inward-open state with respect to the outward-open state where the N-lobe, TM7-8 and C-lobe had mean RMSDs of 1.49±0.17 Å, 2.08±0.43 Å and 2.20±0.16 Å, respectively. In conjunction, these results support the rocker-switch model that has been proposed for DHA2 efflux pumps and indicate conformational stabilization is dependent on specific residue interactions.

Polar residues bridge the N- and C-lobes to stabilize the inward-open conformation, involving residues that are conserved across the DHA2 family. These include hydrogen bonding between the conserved Q302 and the backbone carbonyl of M40 (which varies in the family). Here, the presence of a highly conserved proline, P43, kinks the helix of TM1 and facilitates hydrogen bonding with the M40 backbone (Figure S3D). Mutagenesis data show that substitution of P43 moderately impairs resistance levels (Table 1), suggesting that the proline-induced helical distortion is functionally important. In MD simulations, Q302 interacts with M40 for over 86% of the total simulation time, highlighting the importance of these conserved residues in conformational stability. Other significant polar interactions include those between the E169 (TM5) carboxylate with the S308 sidechain hydroxyl and F310 backbone amine, which are maintained for 97.5% and 94.2% of the total simulation time, respectively. It is likely that these interactions further stabilize the inward-open state. The D61 (TM2) acidic side chain forms a direct contact (3 Å) with the backbone carbonyl of V415 (TM13) in the crystal structure of inward-open QacA. The repulsive nature of a carbonyl/carboxylate interaction suggests this coordination is electrostatically unfavorable. Functional characterization shows that mutation of D61 moderately impacts QacA-mediated resistance (Table 1), supporting a structural role for D61. This analysis shows how both polar and hydrophobic interactions appear to stabilize the two different conformations.

In all three replicate MD simulations of the inward-open state, two Na^+^ ions spontaneously entered the TM cavity from the intracellular solution at the start of the simulation and diffused to two distinct TM binding sites. One Na^+^ ion bound at the Na1 site which was also identified in the outward-open conformation. A Na^+^ ion occupied the Na1 site for 70% of the combined 1500 ns of simulation. The remaining Na^+^ ion bound at a site referred to as the Na3 site, formed by the backbone carbonyl of V31 (TM1), the carboxylate side chains of E407 and D411 (TM13), and three water molecules (Figure 2B and 2E). Notably, mutational analysis of V31, E407 and D411 led to reduced antimicrobial resistance (Table 1), underscoring the critical role of the Na3 site in QacA pump activity. The Na3 site is a relatively low-occupancy site (54 % of the combined 1500 ns of simulation) and is located one helical turn beneath the Na2 site. The proximity of the outward open Na2 site and the inward open Na3 site indicates that this region of electronegativity is functionally important. On transition to an inward occluded state, any Na^+^ ions occupying the Na1 and Na3 sites are trapped within the binding pocket.

### Lipids enter the QacA cavity in the inward-open state

The QacA structure shows an opening between TM5 and 10 on the intracellular side, starting at the Na1 site, involving S154 (TM5) and D323 (TM10), and extending to the intracellular lipid/water interface. The crystal structure shows non-protein density at this opening where a lipid can be modelled, along with a neighboring detergent molecule (Figure S1). In simulations, this TM5/10 opening created a membrane-accessible cleft connecting the inner leaflet and the central binding pocket. During the first 43 ns of all three replicate simulations, the TM5/10 cleft narrows and 1-palmitoyl-1-vaccenoyl-phosphatidylethanolamine (PVPE) lipids from the intracellular leaflet diffuse through the TM5/TM10 cleft and into the binding pocket (Figure 4). In two of the three 500 ns replicate simulations, the lipid acyl tails enter the cavity and occlud the permeation of water. In the remaining replicate simulation, a phosphatidylethanolamine (PE) lipid headgroup enters the TM5/10 cleft after 250 ns of simulation, and moves into the binding pocket, where the cationic amine headgroup of the PE lipid eventually forms a water-mediated interaction with E393 (TM12), which persists for the remaining 20 ns of simulation. Throughout the entire simulation, the PE acyl tails occupy the TM5/10 cleft and do not enter the pocket. In all three replicate simulations, irrespective of whether a lipid headgroup enters the binding pocket, the pocket remains occluded by W149, and the PE lipid occupies the region between W149 and the intracellular solvent, preventing the permeation of water (Figure 4, Figure S3A-B).

**Figure 4.**
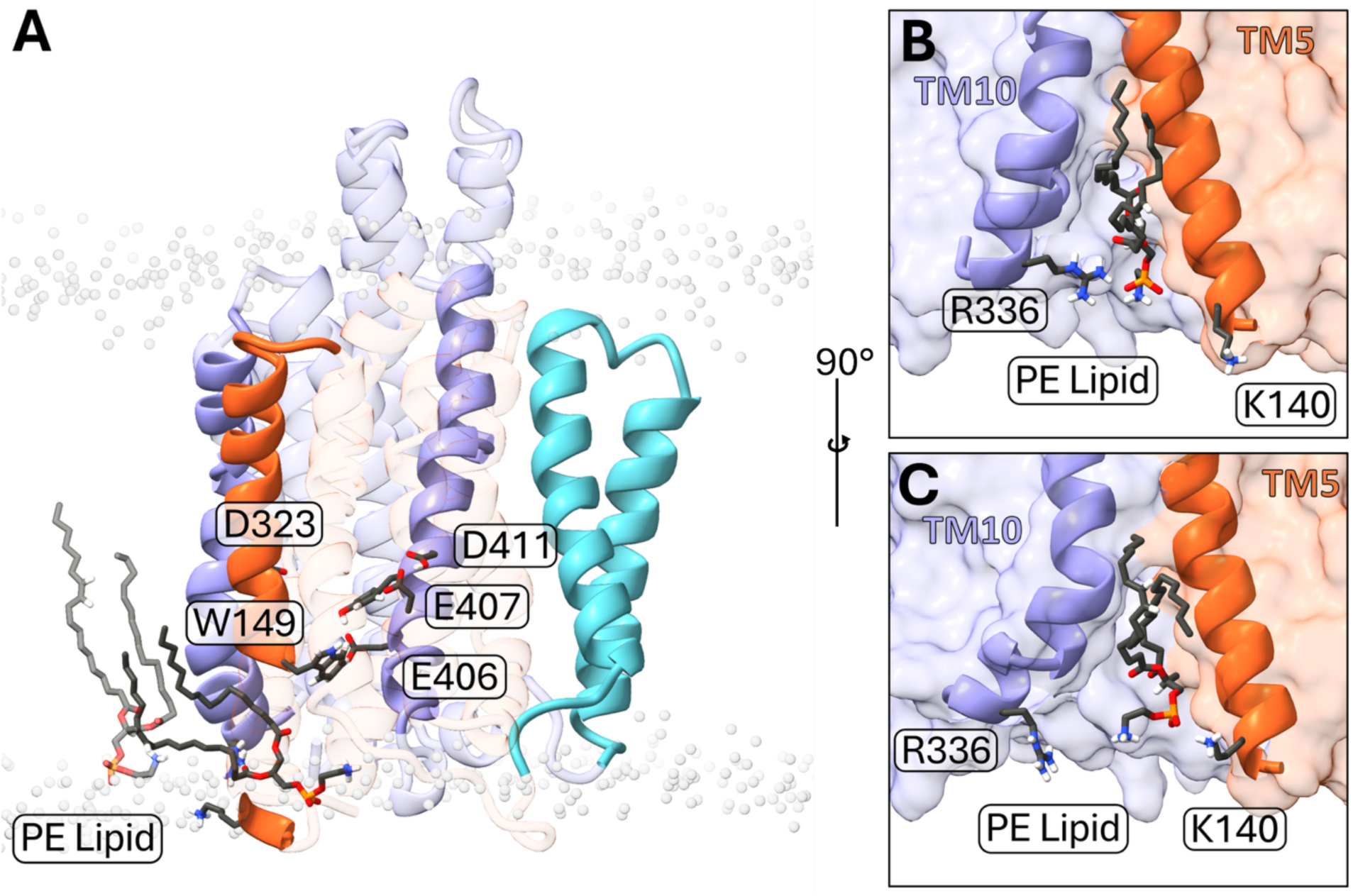
Spontaneous translocation of a membrane lipid into the cytosolic opening of the inward-open QacA binding cavity. A: A PVPE (1-palmitoyl-1-vaccenoyl-phosphatidylethanolamine) lipid from the inner leaflet of the membrane entering through the TM5/TM10 opening of inward-open QacA, below the central binding pocket. B-C: The translocation of the lipid is facilitated by the ‘paddle’ like motions of R336 and K140. Membrane Phosphate headgroups directly adjacent to QacA are represented as grey circles.

The two additional helices found in DHA2 family transporters (TM7 and TM8) form a separate structural domain that reorient relative to both N- or the C-lobes during the structural transition from outward to inward-open, involving different interfaces between TM7 and either TM2 or TM13. In the outward-open state, TM7 interacts mostly with TM2 through a set of hydrophobic contacts but also, notably, via an ionic interaction between K228 (TM7) and D61 (TM2), both residues being conserved within close orthologs of QacA, but not throughout the DHA2 family. MD studies of the outward-open state also reveal a persistent polar interaction between the side-chain hydroxyls of S231 (TM7) and S423 (TM13). In the inward-open state, TM7 reorients upwards and outwards relative to TM2, and the interface is reorganized, with the ionic bond relocating on the other side of D61 (Figure S3E-H). This could suggest a role for the TM7-TM8 domain in the structural transition as supported by the lack of transport activity observed upon deletion of the two helices [20]. MD of the inward-open state shows a persistent salt bridge between K228 (TM7) and D61 (TM2), for the entire 1.5 µs simulation, as well as a relative shift of TM7 extracellularly of TM13 positioning the hydroxyl of S423 (TM13) between the hydroxyl of S231 (TM7) and the backbone carbonyl of I227 (TM7).

There is a hydrophobic protein-protein interface between the accessory TM7/8 helices and TM2 and TM13 from the two main 6-helix bundles. Within the extracellular membrane leaflet, these helices form a compact protein-protein interface, but within the intracellular leaflet, the helices were loosely associated forming a hydrophobic cavity between the helices. This cavity was lined by residues A68, I72 (TM2), M220, L223 (TM7), I251 (TM8), L266 (TM8-9 Loop), A404, V405, L412 (TM13) (Figure S6A-D). In all three 500 ns replicate simulations of the outward open state, the acyl tails of the neighboring lipid from the inner leaflet entered this hydrophobic region at the start of the simulation and remained there for the entire 500 ns simulation. The negatively charged phosphate of the lipid formed a salt bridge with K256 (TM8) and K272 (TM9), stabilizing lipid binding. This lipid binding mechanism was observed in all simulations done throughout this study and occurred with the lipids phosphatidylglycerol (PG), PE and cardiolipin.

### A substrate-bound structure reveals conformational adaptation of the binding pocket

To understand substrate binding by QacA, crystallization screenings were performed in the presence of various known QacA substrates. Incubation with ethidium bromide led to the formation of crystals diffracting to about 2.8 Å in the best direction, revealing a different space group than the apo crystals. This ethidium-bound structure was solved by molecular replacement using the apo outward-open conformation for the search model. The asymmetric unit contains two QacA molecules in the outward-open conformation and structure refinement revealed an additional density in one of them, where a molecule of ethidium can be modelled. As shown in Figure 5A and B, the ligand binds in the center of the TM bundle, sandwiched between the N- and the C-lobe and is exposed to the solvent on the extracellular side (Figure 5C). On the C-lobe, E407 and D411 (TM13) directly coordinate the ligand by interacting with the amine of the aniline groups (Figure 5D and E). Both residues are key for the function of QacA as mutation leads to loss of resistance to ethidium (Table 1).

**Figure 5.**
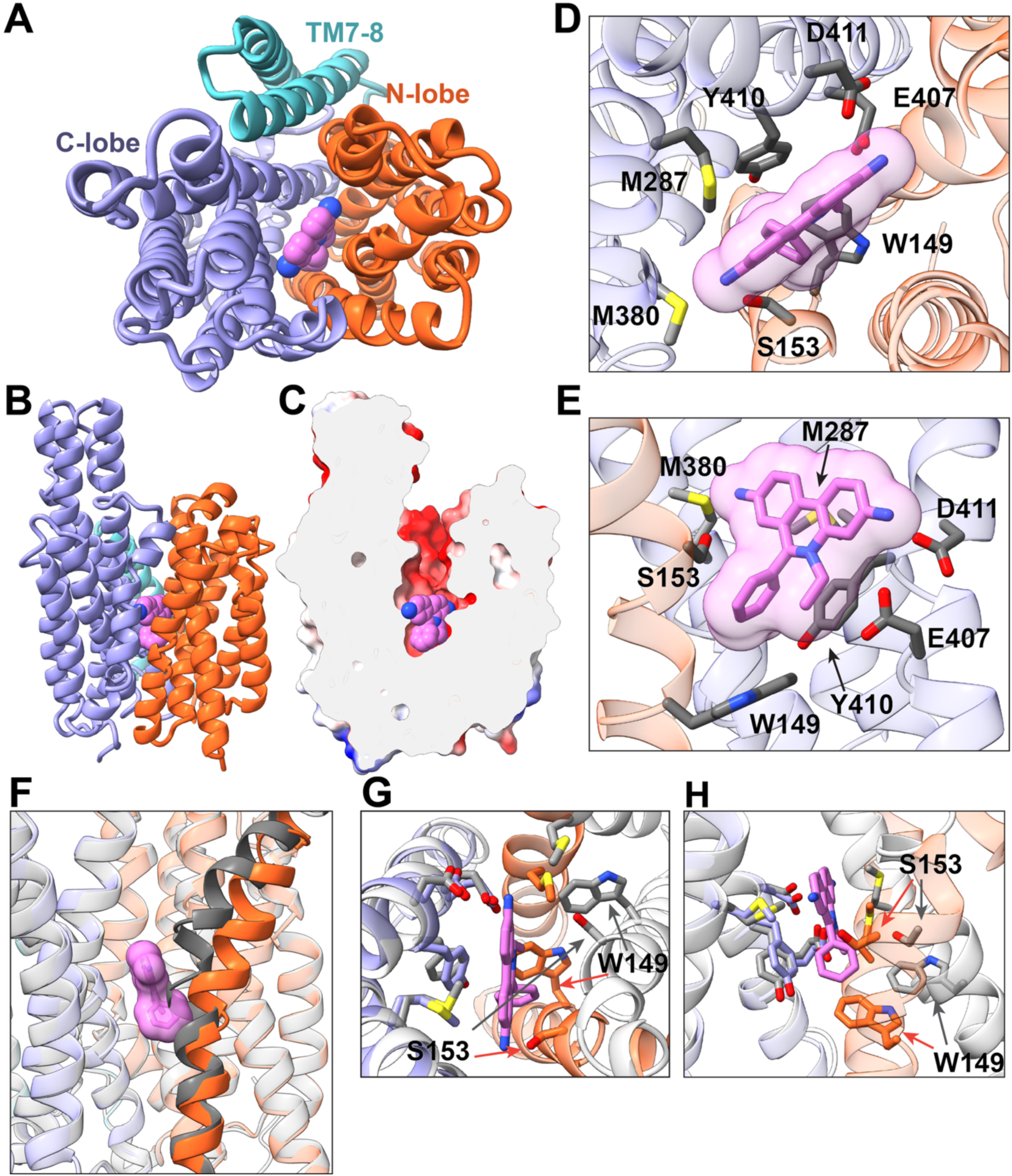
Crystal structure of ethidium-bound QacA. A-B: top and side views of the ethidium-bound QacA showing the overall location of the binding site. C: The molecular surface of QacA was colored by electrostatic charge distribution (using ChimeraX) and cut in the middle to highlight the outward-open state. D-E: Top and side detailed views of the binding site and interacting residues. The molecular volume of ethidium is shown in semi-transparent surface. F: Comparison of the position of TM5 in the ethidium-bound (orange) and ligand-free (grey) states showing that, in the apo state, TM5 overlap with the observed position of ethidium. G-H: top and side views comparing the position of the binding residues in the bound outward-open state (colored) and inward-open state (white) showing that binding side formation requires transition to the outward-open conformation

Compared to the apo-state, Y410 has reoriented away from the N-lobe and points down towards E406 and S387 with its ring parallel to the aromatic cycles of ethidium and close enough (<3.5 Å) to contribute to ligand binding through pi-pi interactions. Binding to the N-lobe involves the second aniline nitrogen of ethidium which appears to point towards S153 in TM5, suggesting a direct interaction. This was confirmed by the reduced resistance to ethidium exhibited by the QacA S153C mutant (Table 1). One helical turn below, W149 also coordinates interactions with ethidium, through possible pi-pi interactions with the benzene ring. Interestingly, a W149A mutation led to decreased ethidium bromide transport but not a W149F change [21] indicating that aromatic interactions are important in the transport of ethidium bromide.

The binding pocket of QacA contains several methionine residues which likely play a role in coordinating ligands. In the ethidium-bound structure, the sulfur atom of M380 is located close (<3 Å) to the nitrogen of the second aniline, likely forming a weak hydrogen bond (Figure 5D). With its sulfur atom located less than 5 Å from one of the aniline rings, M287 is ideally oriented to interact with ethidium through thioester-aromatic interactions, as supported by the fact that its mutation to either cysteine or alanine leads to a strong decrease in ethidium resistance (Table 1). Although often overlooked, thioester-aromatic interactions are known to be important for protein-ligand contacts [22]. The presence of several methionine residues within the binding pocket, which bear sulfur atoms in their flexible sidechain, could contribute to the versatility of the binding pocket in accommodating structurally diverse ligands as those known to be transported by QacA.

Therefore, coordination of ethidium between the two lobes involves residues on TM9 (M287), TM12 (M380) and TM14 (E407, Y410, D411) on the C-lobe and TM5 (W149 and S153) on the N-lobe. This binding site is strictly enabled by a local rearrangement of TM5. Indeed, as shown in Figure 5F, the center of TM5 is moved by about 4Å compared to the apo structure, thus accommodating the presence of the substrate as the position of helix in the apo state (dark grey ribbon in Figure 5) would directly clash with the benzene ring of ethidium. The motion includes local rotation of the helix, moving S153 away from W149 (TM5) and enabling its interaction with the aniline of ethidium.

Remarkably, structural comparison reveals no other conformational difference between the apo outward-open crystal structure and the ethidium-bound structure (RMSD < 0.7Å). This suggests that the conformational flexibility of TM5 underpins the structural adaptation of the TM bundle required to accommodate ligand(s). This hypothesis was investigated by following backbone dynamics using Hydrogen-Deuterium exchange coupled to mass spectrometry (HDX-MS) on detergent-solubilized QacA in the absence or presence of ethidium bromide (Figure S7). This revealed that deuterium uptake is on average higher in TM5 than other helices (except TM10, which is neighboring TM5 in the structure), supporting the above hypothesis that this region is flexible to accommodate multidrug binding. Interestingly, incubation with ethidium bromide led to an increase in H/D exchange in various parts of the protein, suggests that substrate binding augments conformational dynamics (Figure S7).

To examine ethidium binding to QacA, triplicate 500 ns MD simulations were initiated using the crystallographic coordinates of the ethidium-bound conformation for the outward-open state while molecular docking was used for the inward-open conformation. In the outward-open QacA conformation, ethidium remained stably bound in its crystallographic binding pose throughout all three replicate simulations, with an all-atom RMSD of 0.65 ± 0.25 Å across the combined simulations. The aniline amine closest to the quaternary ammonium of ethidium interacted with E407 for over 90 % of simulation and the opposing aniline amine (closest to the phenyl group) interacted with D323 for over 84 % of the simulation (Figure 6A-C, Table S2). Interestingly, geometric constraints of the central binding pocket prevent direct coordination of the quaternary ammonium in ethidium by acidic residues (Figure 6), suggesting the quaternary ammonium in ethidium may contribute primarily to the selectivity, uptake and permeation of substrates, rather than its binding. The stability of ethidium in the crystallographic binding site, and the long-timescale interactions with the acidic E407 and D323 residues, indicate the presence of a single stable binding site in outward-open QacA that is primed for substrate release and efflux.

**Figure 6.**
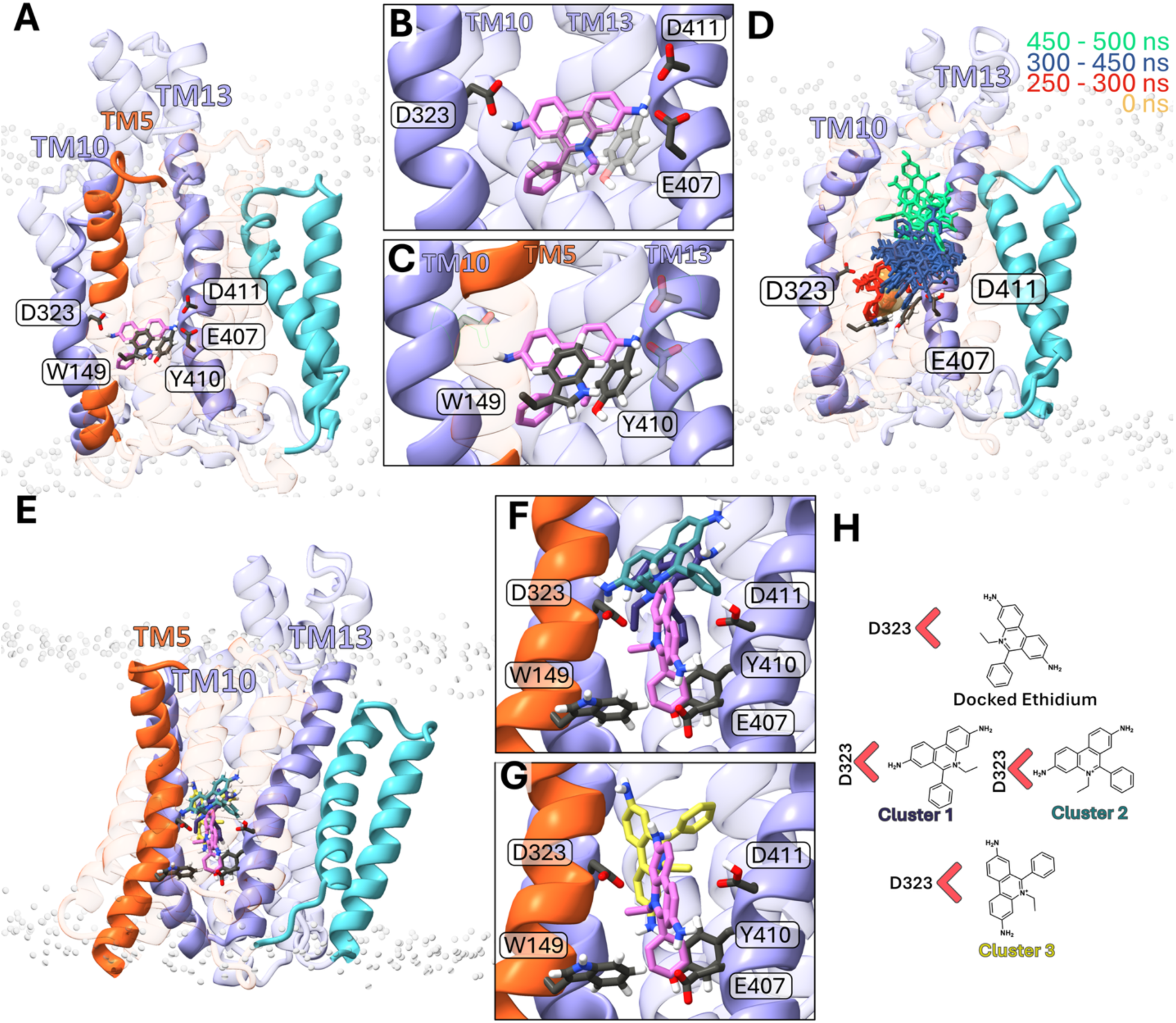
Ethidium binding outward-open and inward-open QacA. A: stable binding pose of ethidium within outward open QacA and interacting B: polar and C: hydrophobic residues with proposed proton switch, E407, deprotonated. D: Overlay of ethidium orientations (every 10 ns) adopted throughout the final 250 ns of one Molecular Dynamics simulation replicate where ethidium left the crystallographic binding site (highlighted in orange). E: Initial docked (pink) and most populated clusters F: purple, blue and G) yellow of ethidium in inward-open with H: respective orientations within the binding pocket.

QacA is a proton antiporter [18], where proton transfer takes place via protonation/deprotonation of charged residues along a solvent-accessible pathway [23]. Calculation of the pKa of the ionizable residues of QacA using PROPKA [24] indicates the pKa of E407 is increased to 12.34, indicating E407 is protonated in the outward-open ethidium bound state. In simulations with E407 protonated, ethidium binding is destabilized and leaves its crystallographic binding site in two of the three 500 ns replicate simulations moving towards the extracellular entrance of the central binding cavity. Destabilization of ethidium binding with the protonation of a key binding residue suggests that this is a likely step in the efflux mechanism of QacA. In the third 500 ns simulation, the ethidium aniline group interacted with the M287 sidechain methyl. Mutations of M35 or M287 decreases resistance to ethidium and other substrates (Table 1), supporting the importance of these thioester-aromatic interactions in substrate efflux [25].

In contrast to simulations of the outward-open conformation, ethidium rapidly left its docked location within the binding site in simulations initiated from the inward-open conformation and diffused toward the extracellular side of the central binding pocket (Figure 6). In each location, one of the two opposing aniline amine groups of ethidium formed electrostatic interactions with either D323 from the Na1 binding site (one replicate), or E407 (remaining two replicates) (Figure 6, Table S2), while Na^+^ ions rapidly entered the TM cavity and bound to the Na1 and Na3 sites. It should be noted, that while some residues are in contact with ethidium for more than 50% of the cumulative simulation time, there is no one clear binding site identified by MD simulation.

## Discussion

This work solves, for the first time, the high-resolution structures of biologically relevant states of QacA, a multidrug transporter implicated in drug resistance within clinical strains of *S. aureus*. Recent advancements in structural biology have enabled the resolution of multiple protein conformations from a single sample, such as the occluded conformations of NorA, which have helped characterize its conformational landscape [26].

The ethidium-bound structure presented here offers novel light on multidrug binding by efflux pumps. For most families of bacterial efflux pumps, only a small number of structures bound to substrates have been obtained at high enough resolution to clearly describe the ligand coordination. Seminal work on the RND protein-family members *E. coli* AcrB has shown that it can coordinate diverse ligands mostly through hydrophobic and aromatic interactions [27, 28]. On the other hand, recent work on *A. baumannii* AdeB shows coordination of ethidium also involves aromatic stacking, polar interactions (including between a glutamate and one of the aniline groups) and pi-thioester interaction with a methionine [29].

While aromatic and polar side chains have been often described as important for binding heterocyclic groups (often found in antimicrobials), the presence of six methionine residues in the binding cavity between the N- and C-lobes of QacA supports a role in coordination of ligands with different aromatic structures. Indeed, while the long aliphatic side chain of methionine can provide an adequate apolar environment for hydrophobic substrates, its sulfur atom can interact with the pi-electron systems of aromatic rings to contribute to the stability of the protein-ligands interaction [22]. Considering the conformational flexibility of their long-side chain, methionine residues could therefore provide versatile systems to coordinate structurally diverse molecules with aromatic groups.

Interestingly, detailed analysis of the different structures of ethidium-bound AdeB, showed slight differences in the binding poses of ethidium (up to 2 Å RMSD) [29]. In agreement with the idea of a versatile coordination (where one pocket can bind several ligands) this shows that a single ligand can bind with some variability to a multidrug transporter and rationalizes the changes in binding poses observed between the crystal structure of ethidium-bound QacA and MD simulations.

A remarkable feature of the crystal structures is the local deformation in TM5 in the ethidium-bound state which appears to be required to sterically accommodate the ligand and enabling its coordination by W149 and S153 (TM5). This conformational change between TM5 of the apo and the ethidium-bound state not only displaces the center of the helix away from the protein center but also involves local rotation between residues 150 to 160 (while the ends of the helix are almost identical). This leads to relocation of S153 (which would otherwise overlap with ethidium) enabling its interaction with the nitrogen of the aniline. The local rearrangement of the helix extends up to the start of motif C [16], which is found in most MFS antiporters and is known to be important for structural and functional integrity of these transporters as mutations of conserved residues typically leads to impaired transport [30], including for QacA [31]. Although with smaller amplitude than for QacA, local structural changes in the center of TM5 have been linked to substrate binding of another multidrug MFS, MdfA [32], suggesting a possible coupling between substrate binding and global conformation changes (i.e. through motif C). This suggests that adaptation to structurally diverse ligands require local plasticity of the binding pocket, including deformation of helices.

By combining experimental and computational data, we can propose a general model for ethidium bromide transport by QacA to be formulated (Figure S8). In the resting state (step I in Figure S8), the extracellular side of the transporter is expected to be closed most of the time to prevent futile proton entry while the intracellular could be either open (inward-open) or also closed (adopting an occluded state that we did not capture by crystallography) with possible conformational exchanges between these states. As the MD simulations show that, in the inward-open conformation, lipids from the inner leaflet of the membrane bilayer spontaneously enter the TM bundle between TM5 and TM10, this interface would provide a direct entry pathway for membrane-embedded drugs to access the binding pocket (step II, Figure 8S). This is in line with early kinetics studies which indicate that multidrug transporters, like QacA, likely capture drugs from the inner leaflet of the membrane [18, 33]. According to our simulations, binding of ethidium in the inward-open state is non-specific, in the absence of structural change the substrate will eventually leave the pocket and move back to the membrane without being expelled. Therefore, we propose that high affinity binding would happen upon a spontaneous (albeit transient) transition to an inward-closed state (*step III*, Figure S8) which would bring the intracellular parts of TM4 and TM5 closer to TM9 and TM13, thus forming the binding site as observed in the crystal structure and simulations. We propose that the substrate-stabilized closing of the intracellular side leads to a (either concomitant or subsequent) opening of the extracellular side, leading the ethidium-bound conformation observed by crystallography (*step IV*, Figure S8). This would agree with previous work on the MFS multidrug transporter LmrP, where it was shown that substrate binding stabilized the outward-open state for a variety of ligands. Importantly, here MD simulations indicate that stable binding of ethidium requires a deprotonated E407 while, in the ligand-free outward-open state, this residue is expected to be protonated. This provides a simple mechanism for extrusion as, the outward-open conformation would enable solvent/proton entrance and subsequent protonation of key side chains (such as E407) leading to substrate-release (*step V*, Figure S8). Protonation of E407 has previously been deemed essential for export by QacA [20]. Therefore, the decrease in binding affinity is not, *per se*, due to a conformational change but rather to modification of the electrostatics of the binding pocket upon protonation, providing a general model to explain extrusion of structurally diverse substrates by a single transporter. Finally, the bound proton is translocated through the transporter and released into the cytoplasm, consistent with an antiport mechanism, allowing the transporter to reset to the resting state (*step VI*, Figure S8), through a molecular mechanism that remains to be elucidated.

## MATERIALS AND METHODS

### Bacterial strains, plasmids, growth conditions and mutagenesis

Two *E. coli* strains were used in this study. Strain DH5α [34] was employed for site-directed mutagenesis, plasmid propagation and antimicrobial susceptibility assays and a BW 25113 ΔAcrAB derivative [35] for QacA overexpression. Bacterial cultures were routinely grown in Luria-Bertani (LB) or Terrific Broth (TB) medium at 37°C with 100 μg/mL ampicillin where appropriate.

For QacA overexpression, an overnight culture of a pBAD-derived construct (QacA-ccHis) was used to inoculate TB medium (1:50 dilution) and grown to OD₆₀₀ ≈ 0.9. Expression was induced with 0.02% (w/v) L-arabinose, followed by incubation for 4 h at 30°C. Cells were harvested by centrifugation at 7000 × g for 15 min at 4°C and stored at -80°C in HEPES buffer.

Site-directed mutations in the *qacA* gene were generated using the plasmid pSK7201, a pBluescript II SK-based vector encoding C-terminal 6xHis-tagged QacA, codon optimized for expression in *E. coli* [36]. Mutations were introduced with complementary oligonucleotide primers [17] and all constructs were verified by DNA sequencing (Australian Genomic Research Facility, Adelaide) to confirm the presence of the desired mutations and the absence of unintended changes within the full *qacA* coding sequence.

Nanobodies were cloned in the pXAP100 vector. pXAP100 is similar to pMES4 (Genbank GQ907248) but contains a C-terminal His6-cMyc tag that allows cloning of the VHH repertoire via SfiI-BstEII restriction sites.

### Antimicrobial susceptibility testing

To assess the antimicrobial resistance activity of QacA variants, MIC analyses of various antimicrobials were conducted using the agar dilution method as described previously [21]. Briefly, midlog-phase bacterial cultures of *E. coli* DH5α carrying WT QacA or its mutants were spotted onto agar media. Mueller-Hinton agar plates were prepared with a gradient of antimicrobial concentrations, including: ethidium bromide (25 to 450 mg/L, in 25 mg/L increments), rhodamine 6G (200 to 1,200 mg/L, in 50 mg/L increments), benzalkonium chloride (20 to 150 mg/L, in 10 mg/L increments), dequalinium chloride (50 to 450 mg/L, in 25 mg/L increments), chlorhexidine dihydrochloride (0.5 to 12 mg/L, in 0.5 mg/L increments) and pentamidine isethionate (60 to 400 mg/L, in 20 mg/L increments). The plates were incubated at 37°C for 24–48h. The MIC was determined as the lowest concentration of antimicrobial agent that completely inhibited visible bacterial growth.

### Preparation of membrane vesicles and QacA expression analysis

Frozen cells were thawed and supplemented with Complete protease inhibitors (Roche). Samples were homogenized using a high-pressure homogenizer (12,000 psi, 3–6 passes). Cell debris and unbroken cells were subsequently removed through centrifugation at 17,000xg, 4 °C, for 30 minutes. Membrane vesicles were then isolated by ultracentrifugation at 125,000×g for 2 hours at 4 °C, resuspended in high-salt buffer (100 mM HEPES, pH 7.0, 600 mM NaCl, 20% glycerol), flash-frozen in liquid nitrogen, and stored at −80 °C for further use.

Expression levels of QacA and its mutated derivatives were determined as previously described [17, 31]. The total membrane protein concentration was determined using a Protein Assay Reagent Kit (Bio-Rad). Equal amounts of total membrane protein were resolved by SDS-PAGE and transferred to PVDF membranes. QacA protein was detected using an anti-6xHis-tag antibody (Rockland Immunochemicals, 1:5,000 dilution). Blots were imaged using the ChemiDoc™ MP Imaging System (Bio-Rad) and protein band densities were quantified with ImageLab software (Bio-Rad) to assess relative expression levels.

### Purification of QacA

Membrane vesicles were thawed and solubilized with an equal volume of 2.4% (w/v) n-dodecyl-β-D-maltoside (βDDM; Inalco) in water for 2 h at 4°C. Insoluble fragments were removed through centrifugation at 100,000×g for 30 min. Centrifugation supernatant was supplemented with 10 mM imidazole, added to Ni-NTA resin (Qiagen) and incubated under agitation for 2 hours at 4 °C, before washing (10 times volume of resin) with wash buffer (50 mM HEPES, pH7, 300 mM NaCl, 10% (v/v) glycerol and 0.05% βDDM) containing 20 mM imidazole and elution with 3 CV of wash buffer containing 250 mM imidazole at 4 °C. Imidazole was removed with PD10 desalting columns (GE healthcare) and PD10 buffer (50 mM HEPES, pH7, 150 mM NaCl, 10% (v/v) glycerol and 0.05% βDDM). QacA was then further purified through size-exclusion chromatography (SDX 200 10/300 GL increase; GE lifescience) in SEC buffer (20 mM HEPES pH7, 100 mM NaCl, 10% (v/v) glycerol, 0.02% βDDM) and concentrated as desired using spin-concentrators.

### Crystallization of QacA

For crystallization experiments QacA WT was purified as detailed above, but with the α-isomer of DDM (Avanti) used in the wash buffer, PD10 buffer and SEC buffer, and with the His-tag removed and the protein lipidated prior to SEC purification. To achieve this, after buffer exchange the protein was concentrated to 5 mg/mL and incubated with *E. coli* polar lipids (Avanti), solubilized at 15 mg/mL in SEC buffer with 1% αDDM, at a 50:1 lipd:QacA molar ratio. Simultaneously, QacA was also incubated with 6His-MBP-3C protease (at a mass ratio of10:1) at 4 °C overnight, with uncleaved QacA, free His-tag and 3C protease subsequently removed through reverse-phase IMAC. Immediately prior to crystallization trials QacA was concentrated to 5 mg/mL, supplemented with 1 mM of ligand (or nanobody at a 1.2:1 ratio) and centrifuged at 16,000xg, 4 °C for 10 min. Vapor diffusion crystallization trials were established using 0.2 μL sitting drops.

### Data collection, structure solution and refinement

All the data collection for QacA was performed at the Diamond synchrotron on the I04 beamline. Each dataset was processed automatically with autoPROC+STARANISO. Molecular replacement was performed via PHASER and the input models were obtained by AlphaFold prediction. Model corrections and building was performed with Coot [37] and BUSTER [38] was used for all refinement steps (see Table S1).

### Nanobody production, purification and characterization

Nanodiscs-reconstituted QacA was produced using previously established protocols [8] and used for immunization of a llama. Nanobodies were obtained after phage display selection, using established protocols [39]. After two rounds of selection using detergent-solubilized QacA, a set of candidate binders was isolated and classified according to the sequences of the third complementarity determining region (CDR3) before characterization.

Nanobody expression and purification were performed as previously described [39]. Briefly, nanobodies were produced in Escherichia coli BL21(DE3) pLysS cells (Millipore), purified from the periplasmic extract via either Ni-NTA resin (Qiagen) followed by a size exclusion chromatography on a Superdex 200 Increase 10/300 GL (Cytiva) equilibrated in 20mM HEPES pH 7.5, 150mM NaCl, and 10% (w/v) glycerol.

For dose-response ELISA assays, Nunc maxiSorp 96-well plates (ThermoScientific), were coated with 5 µg/ml NeutrAvidin Biotin-Binding protein (ThermoScientific) overnight at 4 °C and blocked 2 h at room temperature (RT) with 4% milk in phosphate-buffered saline (PBS). Each new reagent addition was preceded by three washes with 200 µl of SEC buffer (20 mM HEPES pH7, 100 mM NaCl, 10% (v/v) glycerol, 0.02% βDDM). Then, biotinylated purified QacA proteins at 5 µg/ml were immobilized 30 min at RT followed by 1 h RT incubation with 100 µl various concentrations (0-20 µg/ml) of purified nanobodies. Signal detection was followed using His-tag specific antibody (0.5 µg/ml - Invitrogen) to detect the nanobodies and secondary antibody anti-mouse coupled to horse radish peroxidase (HRP) (0.5 µg/ml – Millipore). 50 µl of 1-Step UltraTMB-ELISA (ThermoScientific) was used as a substrate for the peroxidase and intensity of the reaction was proportional to absorbance measured at 450 nm after addition of 50 µl H_2_SO_4_ at 1M.

### Molecular dynamics simulations

The X-ray structures of both the apo and ethidium-bound outward-open QacA, and the apo inward-open QacA were used to initiate MD simulations. Prior to simulations, alanine residues at 262 and 263 of the inward-open QacA were computationally mutated to S262 and D263 to match the sequence of ethidium bound outward-open QacA. The nanobody was removed from the inward-open structure and ethidium was docked into the substrate binding site with Autodock Vina [40, 41], using the QacA crystallographic substrate binding residues as a guide. PROPKA was used to calculate the protonation state for all ionizable residues in all three QacA structures [42–45] (Table 2). As protonation and deprotonation of the QacA proton relay network is integral in protein conformation and substrate transport, the protonation state of residues D411, E406 and E407, which are believed to form the proton relay network [11, 20], was manually assigned in each simulation. Table 1 gives an overview of the five QacA systems simulated.

**Table 2.** Protonation of QacA ionisable residues within the binding cavity per PROPKA pKa estimation.

|  | SYSTEM 1 | SYSTEM 2 | SYSTEM 3 | SYSTEM 4 | SYSTEM 5 |
| --- | --- | --- | --- | --- | --- |
| RESIDUE | Outward apo | Outward ethidium | Outward ethidium<sup>a</sup> | Inward apo | Inward ethidium<sup>b</sup> |
| D411 | - | - | - | H | H |
| E406 | H | - | - | - | - |
| E407 | H | H | - | - | - |

<sup>a</sup>Ethidium was docked into the inward open apo crystal structure of QacA with Autodock Vina [40, 41].
<sup>b</sup>Alternate protonation state.
<sup>c</sup>- indicates a negatively charged residue sidechain.
<sup>d</sup>H indicates a protonated, neutral residue sidechain.

In each system, QacA was embedded in a phospholipid bilayer containing 75 % 1-palmitoyl-2-vaccenoyl-phosphatidylethanolamine (PVPE), 20% 1-palmitoyl-2-vaccenoyl-phosphatidylglycerol (PVPG) and 5% cardiolipin (CL), with lipid unsaturated at the n-7 position (vaccenic acid) to reflect the bilayer of the *E. coli* expression system. Parameters were obtained from the Automated Topology Builder and Repository [46] for each lipid (PE molid: 832557; hash: 4e820, PG molid: 828852; hash: dd084, and CL molid: 737061; hash: 884c0) and for ethidium (molid: 1181003; hash: ff144). The system was solvated with the simple point-charge water model and 150 mM NaCl [47]. Counter-ions were added to ensure the overall charge neutrality of the system.

All simulations were performed using GROMACS 2023 [48–51] in conjunction with the GROMOS 54a7 united atom forcefield [52], using a 2-fs time step. All simulations were performed under periodic boundary conditions in a rectangular box. The simulations were equilibrated over 5 ns, using a series of 1 ns simulations with progressively decreasing position restraint force constants of 1000 kJ mol^-1^nm^2^, 500 kJ mol^-1^nm^2^, 100 kJ mol^-1^nm^2^, 50 kJ mol^-1^nm^2^, and 10 kJ mol^-1^nm^2^ on the protein. Simulations were run in an NPT ensemble where the temperature was maintained at 300 K using the velocity rescale thermostat with a coupling constant of 0.1 ps and the pressure was maintained at 1 bar using the Berendsen barostat and a coupling constant of 0.5 ps. The pressure coupling was semi-isotropic with an isothermal compressibility of 4.5 x 10^-5^ bar^-1^. Non-covalent interactions were determined via the Verlet scheme with a cut-off of 1.4 nm. Long range electrostatics were treated with a with a reaction-field and a dielectric constant of 61 [53]. The SHAKE algorithm was used for constraining covalent bonds involving hydrogen atoms and SETTLE was used for water molecules. Each system was simulated in triplicate for 500 ns per replicate. Simulations were visualised in VMD 1.9.4a55 [54].

Analysis was performed across the full 500 ns of simulation for each replicate, unless otherwise stated. RMSD of each simulation system was calculated on frames collected at 1 ns intervals using GROMACS tools. The RMSD of each trajectory was calculated with respect to the starting QacA crystallographic structure after performing a least-squares fit. The contact frequency was defined as the number of frames of the combined replicate simulations that the centre of mass of any given atom of a specified molecule (or ion) was within a distance of 0.4 nm from the centre of mass of any atom of a second molecule. Contact frequencies were calculated for ethidium and sodium ions and expressed as a percentage of the total simulation time. Protein secondary structure was calculated using DSSP (Dictionary of Secondary Structure of Proteins) [55].

### Hydrogen Deuterieum eXchange - Mass Spectrometry measurements

HDX-MS experiments were performed on Waters Synapt G2-Si mass spectrometer. QacA samples were prepared at a concentration of 40 to 60 µM using Vivaspin concentrators with 50 kDa MWCO. All samples were pre-incubated on ice for 30 min, either in the absence or presence of ethidium bromide which was added at a 1:30 protein-to-substrate molar ratio.

HDX labeling was initiated by diluting 7 µL of a protein sample into 63 µL of labeling buffer (20 mM HEPES pD7, 100 mM NaCl, 10% (v/v) glycerol,0.02% βDDM, prepared in D₂O). Labeling reactions were performed at 20 °C using a plate heater and allowed to proceed for defined time intervals ranging from 5 min to 1 h. Labeling reactions were quenched by the manual addition of an equal volume of pre-chilled quench buffer (50 mM phosphate buffer, 1.0% formic acid, 0.05% βDDM, pH 2.3). Following quenching, samples were flash-frozen in liquid nitrogen and stored at -80 °C until HDX-MS analysis. Prior to injection, samples were thawed at room temperature, with a reproducible thawing time of approximately 75 s. Samples were digested online on Enzymate BEH Pepsin Column (Waters Corporation) at 200 µl min^−1^ and at 20 °C with a pressure 9 kPSI. Peptic peptides were trapped for 3 min on an Acquity UPLC BEH C18 VanGuard Pre-column (Waters Corporation) at a 200 µl min−1 flow rate in water (0.1% formic acid in HPLC water pH 2.5) before eluted to an Acquity UPLC BEH C18 Column for chromatographic separation. Separation was done with a linear gradient buffer (3–45% gradient of 0.1% formic acid in acetonitrile) at a flow rate of 40 µL.min^-1^. Peptide identification and deuteration uptake analysis was performed on the Synapt G2 in ESI ± MSE mode (Waters Corporation). Leucine enkephalin was applied for mass accuracy correction and sodium formate was used as calibration for the mass spectrometer. MSE data were collected by a 20–45 eV transfer collision energy ramp. The pepsin column was washed between injections using pepsin wash buffer (1.5 M Guanidinium HCl, 4% (v/v) acetonitrile, 0.8% (v/v) formic acid). A cleaning run was performed every three samples to prevent peptide carry-over. All deuterium time points were performed in technical triplicate.

Acquired reference MS^E^ data were analyzed by PLGS (ProteinLynx Global Server 3.0.2, Waters) to identify the peptic peptides, then all the MS^E^ data including reference and deuterated samples were filtered and processed by DynamX v.3.0 (Waters) for deuterium uptake determination with the following filtering parameters: minimum intensity of 1000, minimum and maximum peptide sequence length of 5 and 25, respectively, minimum MS/MS products of 2, minimum products per amino acid up to 0.30, minimum score of 5, and a maximum MH^+^ error threshold of 15 p.p.m. Peptide identification was considered valid only when a peptide was consistently observed in a minimum of three out of five independent reference samples. When further statistical analysis was required, Deuteros 2.0 (v 2.4.2), developed by Lau et al. [56], was used. Differences between two states were first evaluated using the hybrid significance test, which assesses whether observed deuterium uptake differences exceed a threshold defined by the selected confidence level (98% confidence; *α* = 0.02). Statistical significance was subsequently confirmed using Welch’s t-test.

## ACKNOWLEDGEMENTS

This work was supported by National Health and Medical Research Council Ideas grant 2020421 to MHB, MLO and CG, The Hospital Research Fund to MHB and MLO, and by grants from the FRS-FNRS to CG (J0044.17F, T.0057.15F). This research was supported by the Australian Government’s National Collaborative Research Infrastructure Strategy (NCRIS), with access to computational resources provided by the National Computational Infrastructure and the Pawsey Supercomputing Research Centre through the National Computational Merit Allocation Scheme to MLO. C.G. is a senior Research Associate of the FRS-FNRS and a WELRI Investigator. We acknowledge the support and the use of resources of Instruct-ERIC, part of the European Strategy Forum on Research Infrastructures (ESFRI), and the Research Foundation - Flanders (FWO) for their support to the Nanobody discovery. We thank Alison Lundqvist for the technical assistance during Nanobody discovery

## DATA AVAILABILITY

The atomic coordinates and structure factors reportedin this paper have been deposited in the Protein Data Bank (PDB). The accession numbers for the structures reported in this paper are PDB: 29OM (wt QacA_in outward-open conformation, 29ON (wt QacA_bound to nanobody 89 in inward-open conformation and 29PX (wt QacA_bound to ethidium in outward-open conformation).

The sequence of Nb89 has been deposited as SD-43M6 in the single domain repository https://www.nanosaurus.org/

Molecular dynamics starting conformations, mdp files and topologies are available at https://github.com/OMaraLab/QacA_Structural_2026

## Supplementary tables

**Table S1.** Data Collection and Refinement statistics.

|  | **QacA_OO**<br>**(PDB 29OM)** | **QacA_IO**<br>**(PDB 29ON)** | **QacA_Et**<br>**(PDB 29PX)** |
| --- | --- | --- | --- |
| **Data Collection** |  |  |  |
| Beamline | Diamond Light Source. I04 | Diamond Light Source. I04 | Diamond Light Source. I04 |
| Detector Resolution (Å) | 1.97 | 1.97 | 1.97 |
| Wavelength (Å) | 0.09795 | 0.09795 | 0.09795 |
| Space group | P 21 21 21 | C 1 2 1 | P 21 21 2 |
| Cell dimensions a, b, c (Å) | 61.70. 74.21. 279.06 | 188.30. 94.33. 98.31 | 146.28. 169.86. 61.78 |
| Cell dimensions $\alpha$, $\beta$, $\gamma$ (°) | 90.00 90.00 90.00 | 90.00. 97.47. 90.00 | 90.00 90.00 90.00 |
| Resolution (Å) | 69.77-2.82 (3.03-2.82) | 93.35-3.32 (3.74-3.32) | 110.84-2.79 (3.21-2.79) |
| R<sub>p</sub>im | 0.053 (0.585) | 0.134 (0.528) | 0.022 (0.453) |
| Total number of observations | 358687 (19111) | 117024 (5801) | 380115 (18000) |
| Total number unique | 26962 (1348) | 17025 (851) | 14427 (722) |
| Mean(I)/sd(I) | 9.4 (1.58) | 4.9 (1.5) | 14.4 (2.0) |
| Completeness (spherical) (%) | 84.7 (22.8) | 66.9 (11.3) | 36.7 (5.4) |
| Completeness (ellipsoidal) (%) | 92.4 (39.4) | 90.6 (49.9) | 91.1 (71.8) |
| Multiplicity | 13.3 (14.2) | 6.9 (6.8) | 26.3 (24.9) |
| CC1/2 | 0.999 (0.552) | 0.992 (0.625) | 0.998 (0.808) |
| **Refinement** |  |  |  |
| Resolution (Å) | 2.83 | 3.32 | 2.83 |
| R<sub>work</sub>/R<sub>free</sub> | 0.310 / 0.328 | 0.297 / 0.327 | 0.321 / 0.351 |
| No. Reflections (R<sub>free</sub> set) | 1240 (3.93%) | 838 (4.93%) | 746 (1.90%) |
| Total atoms | 8249 | 9333 | 7499 |
| Non-hydrogen atoms | 8249 | 9333 | 7499 |
| Protein atoms | 7531 | 9079 | 7295 |
| Water molecules | 69 | 0 | 0 |
| Wilson B-factor | 86.1 | 81.0 | 95.4 |
| Average B-factor (all atoms) | 86.0 | 82.0 | 42.0 |
| R.m.s.d. bonds (Å) | 0.0101 | 0.0107 | 0.0102 |
| R.m.s.d. angles (°) | 1.16 | 1.18 | 1.15 |
| Clashscore | 5 | 7 | 8 |
| Ramachandran favoured (%) | 98 | 92 | 93 |
| Ramachandran allowed (%) | 2 | 8 | 7 |
| Ramachandran outliers (%) | 0 | 0 | 0 |

Note :Data shown in parentheses are statistics for the highest-resolution shell.

**Table S2.** Residues in contact (within 4 Å) with ethidium >50% of cumulative Molecular Dynamics simulation time (1500 ns).

| Residue | Contact time |  |  |
| --- | --- | --- | --- |
|  | Outward-open |  | Inward-open |
|  | E407<sup>-</sup> | E407 protonated | D411 protonated |
| T32 |  |  | 69.7% |
| M35 |  |  | 91.6% |
| T36 |  |  | 77.5% |
| S64 |  |  | 62.0% |
| L146 | 51.7% |  |  |
| W149 | 95.5% | 62.2% |  |
| S150 | 64.0% | 57.2% |  |
| S153 | 63.4% | 68.0% |  |
| A157 |  |  | 91.1% |
| V158 |  |  | 51.7% |
| M287 | 95.7% | 64.5% | 94.2% |
| M290 |  |  | 70.5% |
| L294 |  |  | 81.0% |
| M319 |  |  | 84.1% |
| D323 | 84.2% | 78.1% | 88.5% |
| M380 | 94.2% | 60.2% |  |
| L383 | 98.8% | 60.6% |  |
| A384 | 66.4% |  |  |
| S387 | 63.2% | 52.1% |  |
| E407 | 90.4% | 60.7% |  |
| Y410 | 91.2% | 68.4% | 96.7% |
| D411 | 69.1% | 72.3% | 68.4% |
| N414 | 56.9% | 71.5% | 92.1% |

## Supplementary Figures

**Figure S1.**
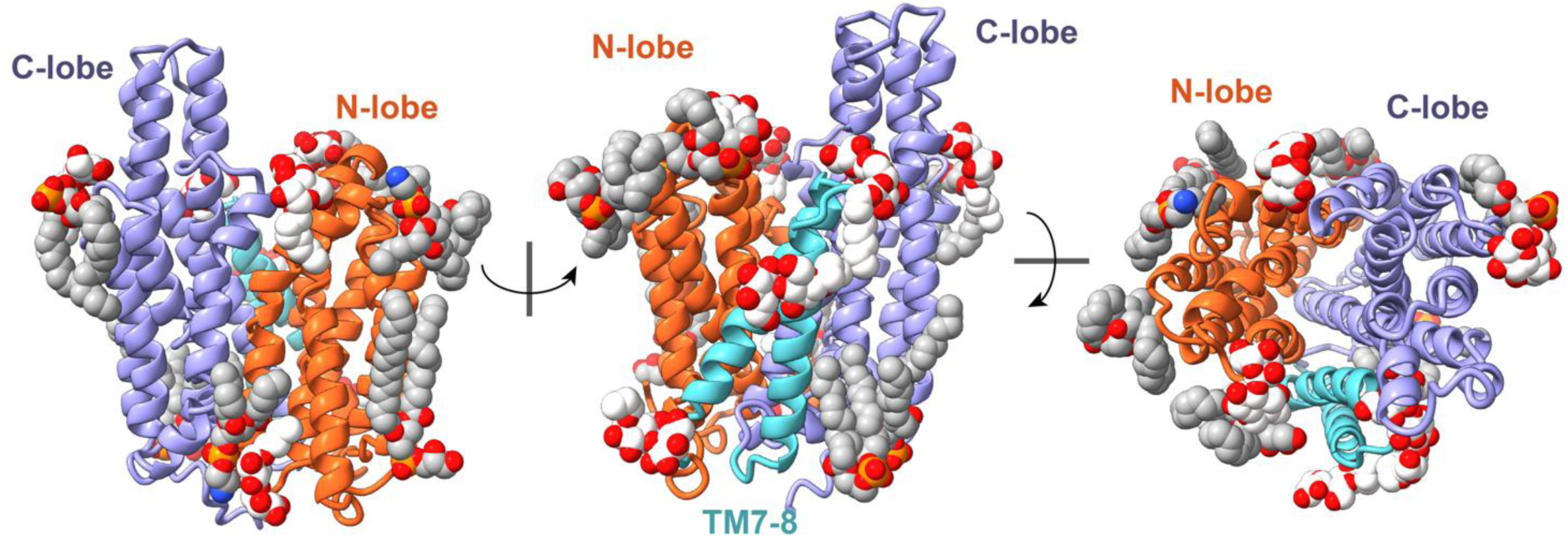
Structure of QacA reveals bound lipids. Ribbon representations of QacA with bound lipids and detergent molecules shown as space-filling spheres (the carbon atoms for lipids and detergent are colored in grey and white, respectively). Lipids identification is tentative.

**Figure S2.**
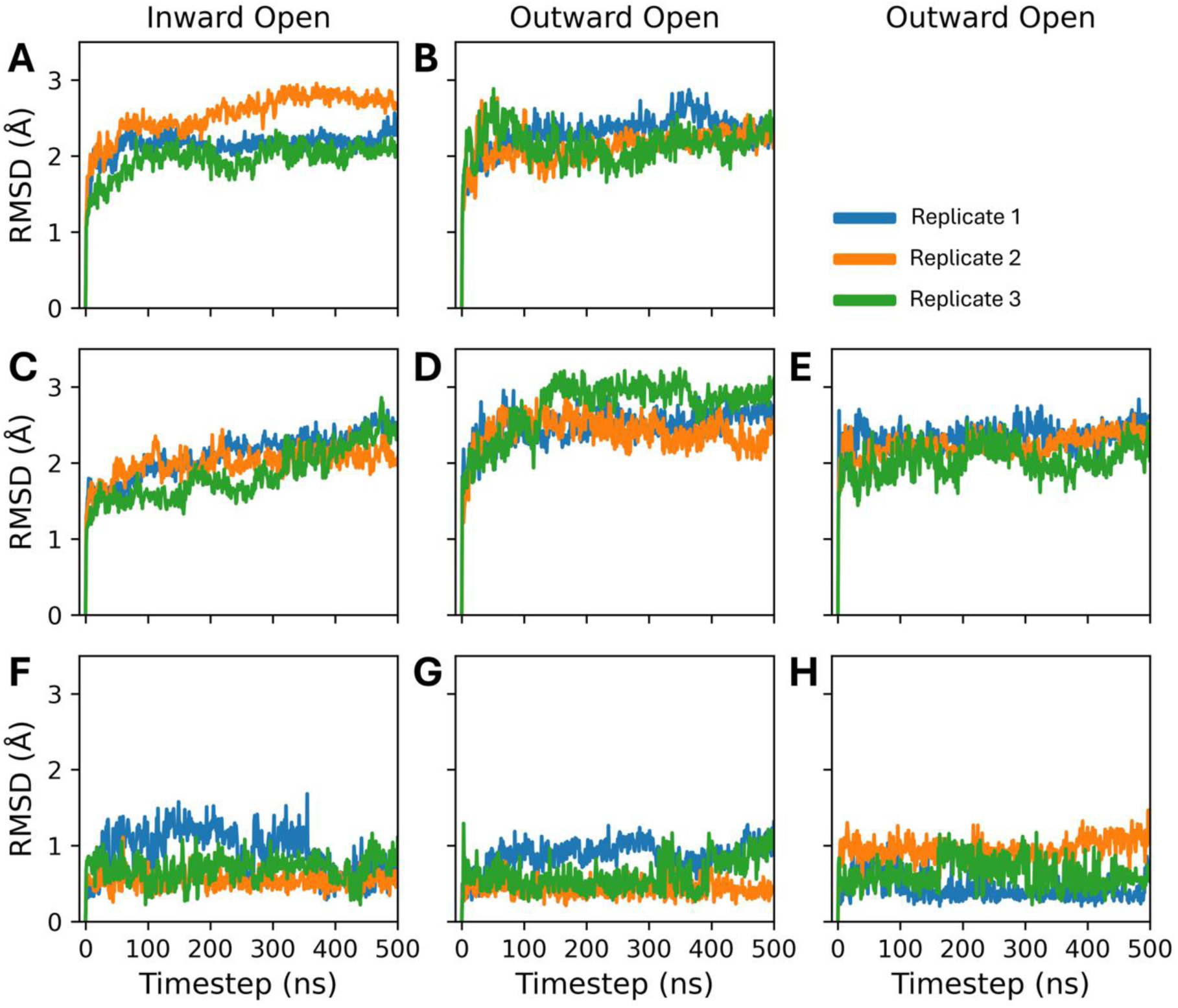
RMSD of QacA backbone and ethidium. Backbone RMSD of A) inward-open apo, B) outward-open apo, C) inward-open ethidium bound, D) outward-open ethidium bound, E) outward-open ethidium bound (E407 protonated) QacA with respect to crystallographic coordinates in triplicate 500 ns Molecular Dynamics simulations. All-atom RMSD of ethidium bound to F) inward-open QacA, G) outward-open QacA and H) outward-open QacA (E407 protonated) with respect to the starting coordinates in triplicate 500 ns Molecular Dynamics simulations.

**Figure S3.**
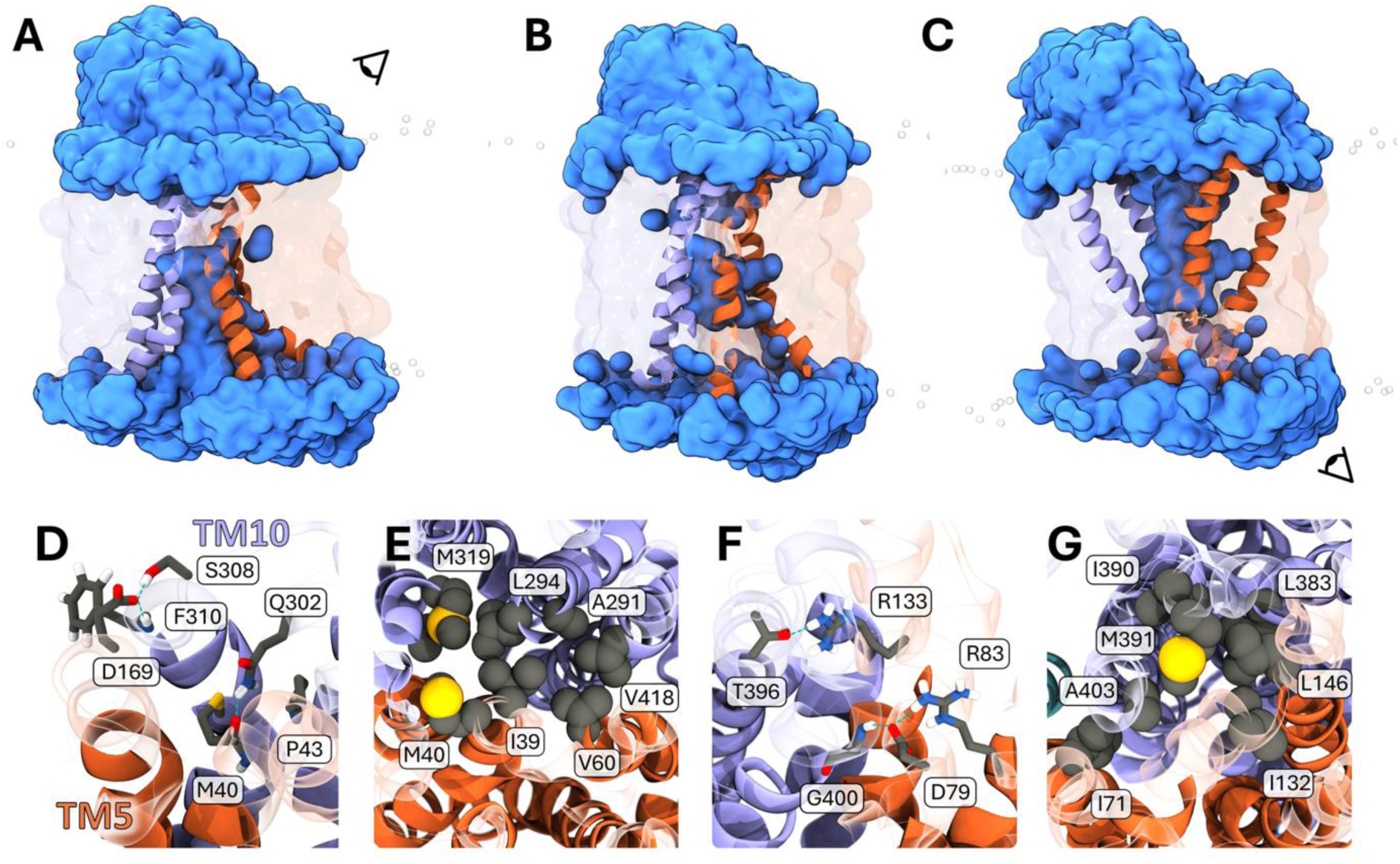
QacA binding pocket solvent accessibility and stabilising polar network of inward-open and outward-open conformation. Solvent occupying the central binding pocket of A) inward-open QacA and C) outward-open QacA. B) The occlusion of cytosolic solvent by a PVPE (1-palmitoyl-1-vaccenoyl-phosphatidylethanolamine) lipid in inward open QacA after 48 ns of simulation time. Solvent (blue) within 10 Å of QacA is shown as a surface representation. D) Polar residues (licorice representation) which stabilise inward-open QacA and E) hydrophobic residues (Van der Waals representation) which prevent solvent penetration to the extracellular space. F) Polar residues (licorice representation) which stabilise outward-open QacA and G) hydrophobic residues (Van der Waals representation) which prevent solvent penetration to the cytosol. Panel A and C have viewing indicators for panel D and F, respectively.

**Figure S4.**
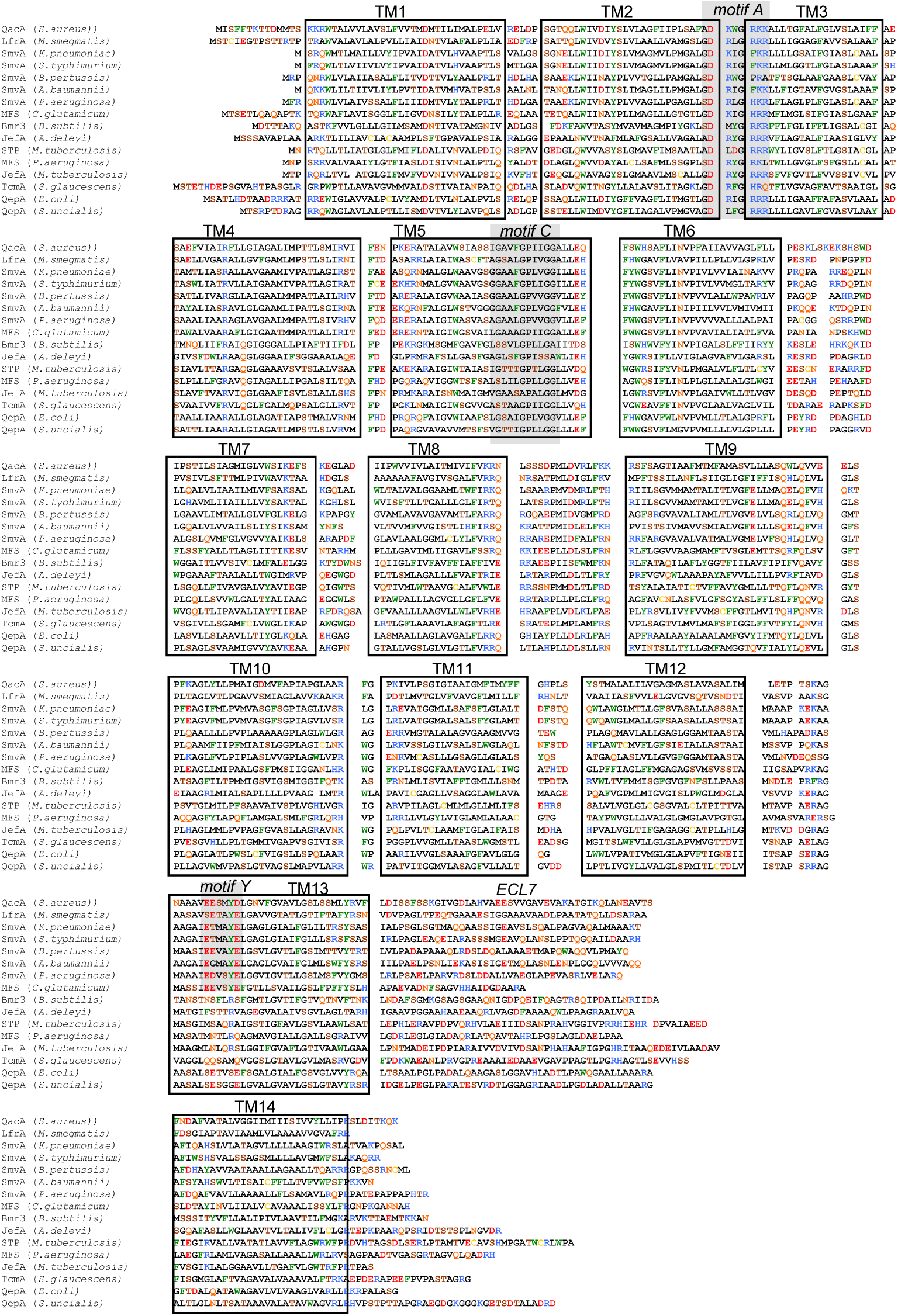
Sequence alignment of QacA and related orthologs. Alignment was based on 3D superimposition (structures and Alphafold models). TM helices and key motifs are highlighted. Uniprot IDs: QacA (*S. aureus*), P0A0J9; LfrA (*M. smegmatis*), A0R5K5; SmvA (*K. pneumoniae*), A0A2H2VR63; SmvA (*S. typhimurium*), P37594; SmvA (*B. pertussis*), A0A381A2V9; SmvA (*A. baumannii*), A0A385EXR3; SmvA (*P. aeruginosa*), A0A9P1VYL7; MFS (*C. glutamicum*), Q8NMG4; Bmr3 (*B. subtilis*), P96712; JefA (*A. deleyi*), A0A6S7A0R1; STP (*M. tuberculosis*), P9WG91; MFS (*P. aeruginosa*), Q9I428; JefA (*M. tuberculosis*), P9WJW9; TcmA (*S. glaucescens*), P39886; QepA (*E. coli*), A5H8A5; QepA (*S. uncialis*), A0A1Q4UZZ7.

**Figure S5.**
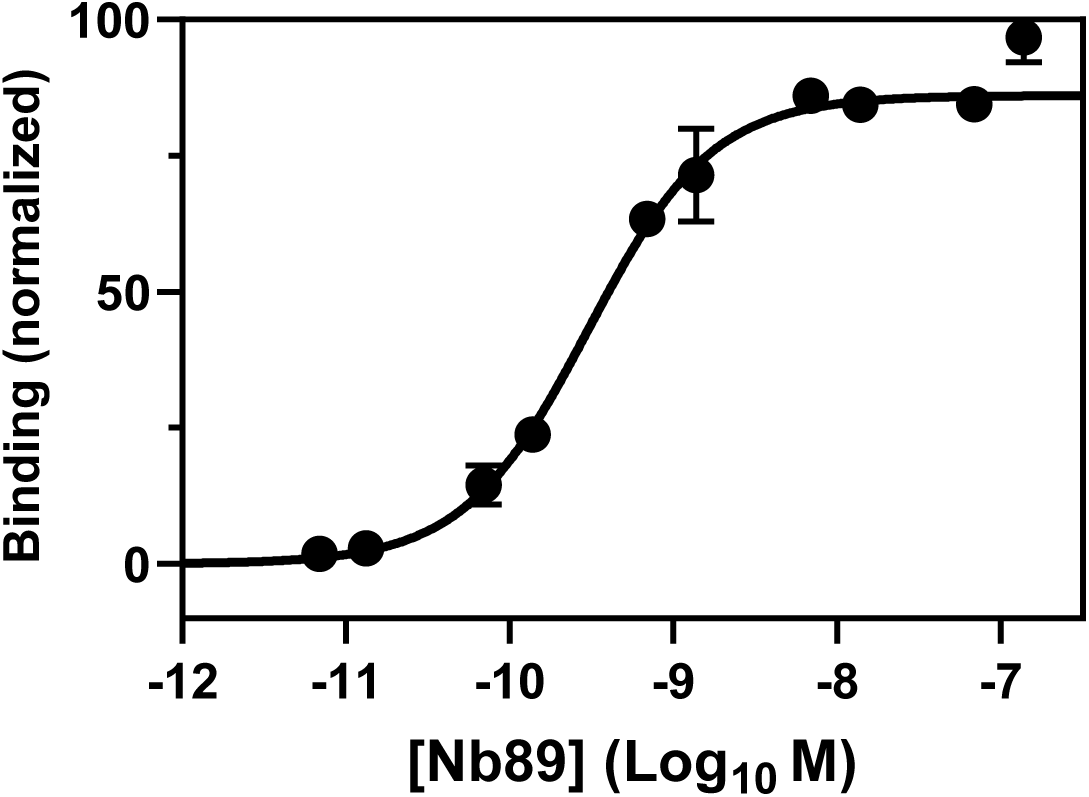
Binding of Nb89 to QacA. ELISA assay using detergent-solubilized, streptavidin-captured QacA incubated with increasing concentrations of Nb89 shows high affinity binding of the nanobody (EC_50_ of ∼3.10^10^M).

**Figure S6.**
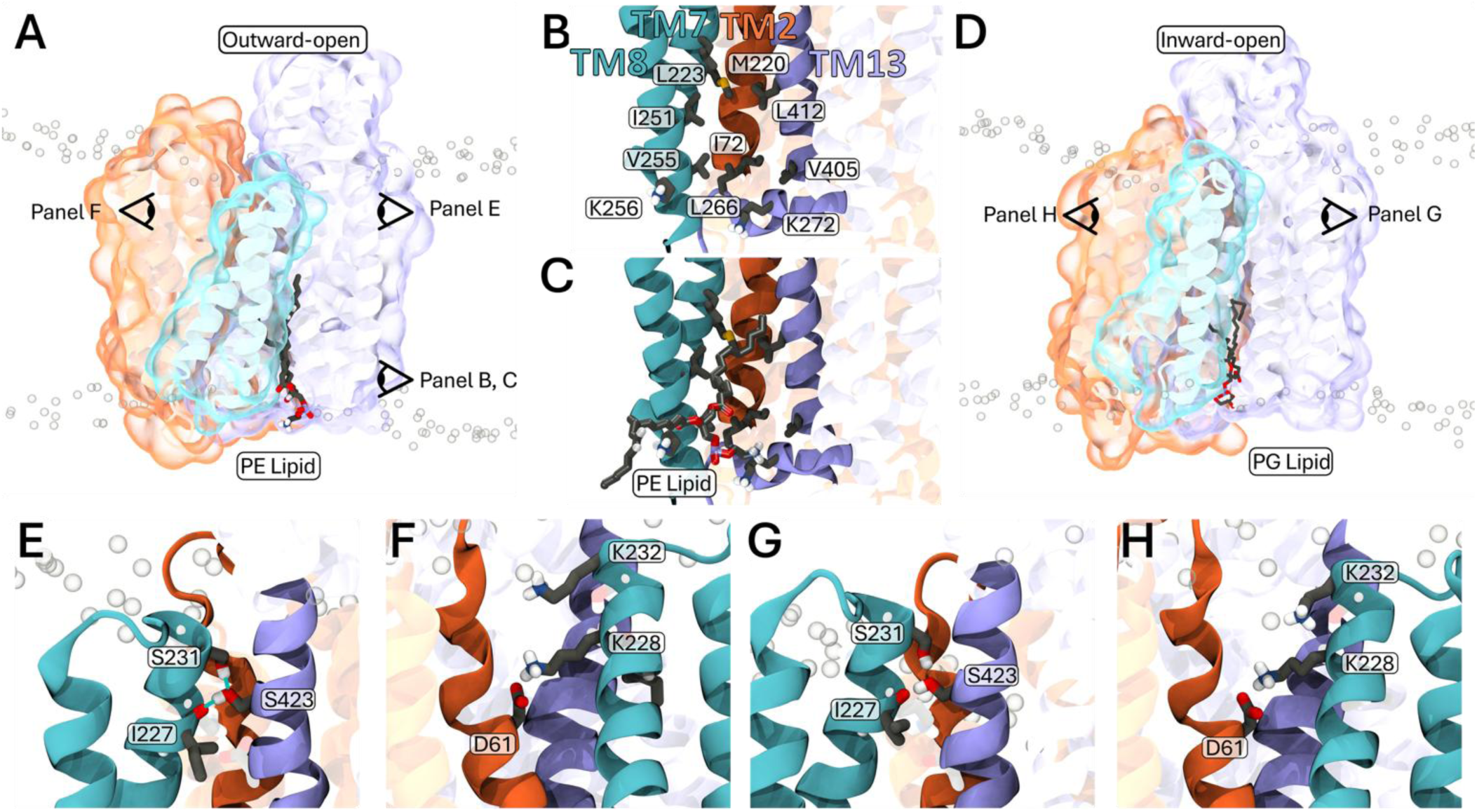
Intracellular leaflet membrane lipids filling a hydrophobic protein-protein cavity adjacent to TM2 (N-Lobe), TM7 & 8 and TM13 (C-Lobe). A) The saturated palmitoyl sn1 tail of a 1-palmitoyl-2-vaccenoyl-phosphatidylethanolamine burying into the hydrophobic cavity formed by TM2, TM7/8 and TM13 in outward-open QacA. B) The hydrophobic residues lining this cavity, with C) the PE lipid depicted. D) The saturated palmitoyl sn1 tail of a 1-palmitoyl-2-vaccenoyl-phosphatidylglycerol lipid entering the hydrophobic cavity, which is still present in inward-open QacA. The N-lobe (orange), TM7/8 (blue) and C-lobe (purple) are shown in cartoon and surface representations, with membrane phosphates a grey circles. Key interactions between TM7/8 and the N- and C-lobes for E-F) outward-open and G-H) inward-open QacA.

**Figure S7:**
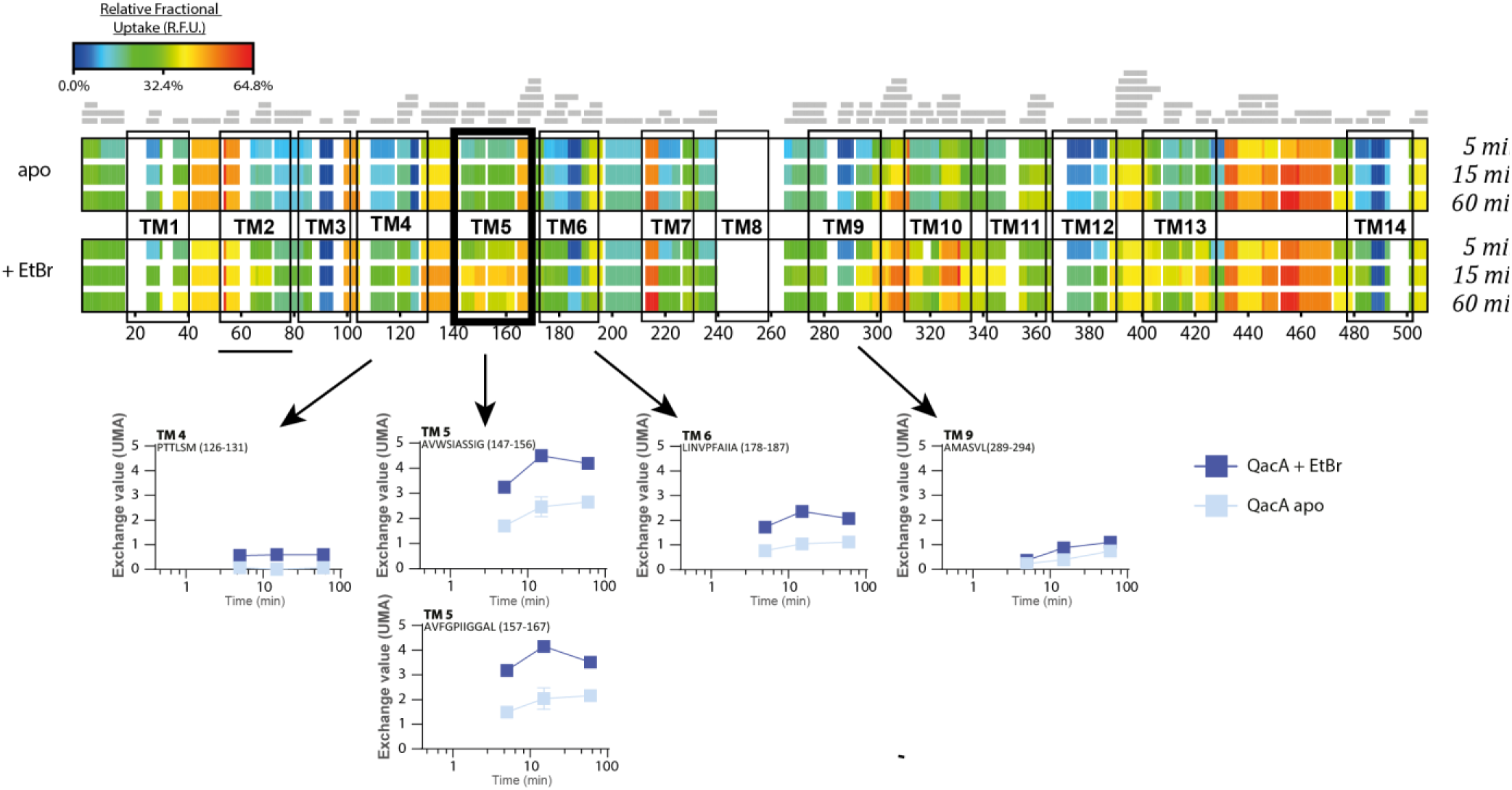
HDX maps of QacA in absence and presence of EtBr. Relative fractional uptake is color-coded from 0% (blue) to 65% (red) at three time points (5, 15 and 60 min) for each peptide, shown along the sequence (numbered below) with TM helices highlighted. Identified peptides are represented by grey bars. Individual graphs show uptake plots for 5 representative peptides from 4 different TM helices. High exchange is observed for TM5, further increased in the presence of EtBr.

**Figure S8:**
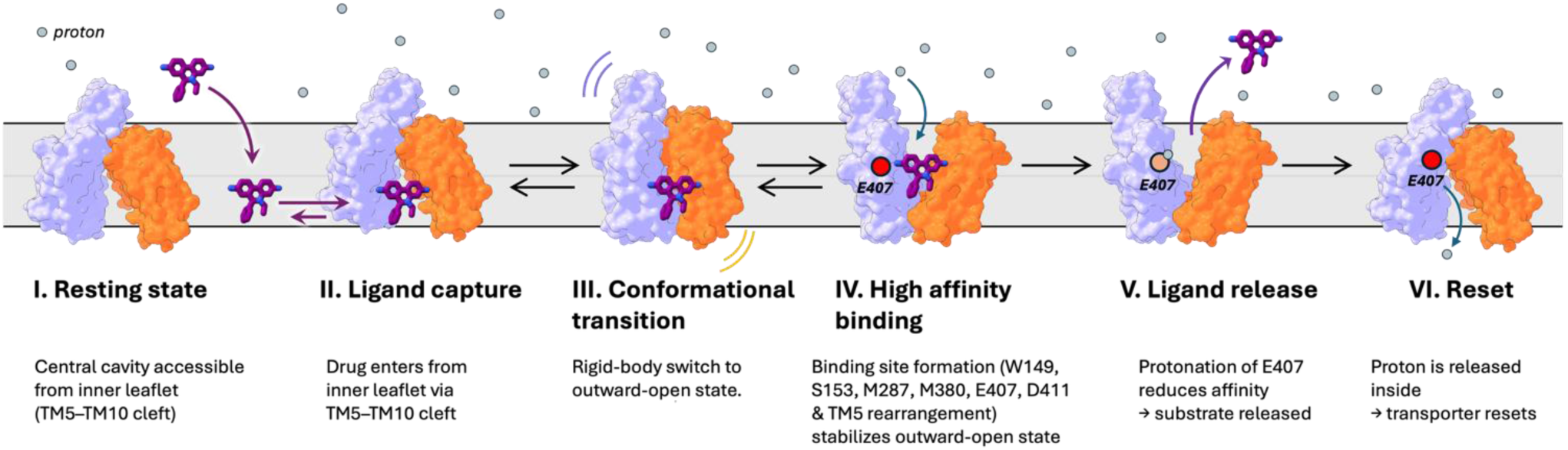
General model for drug extrusion. In the resting state (**step I**), the transporter adopts an inward-open conformation in which the central cavity is accessible from the inner leaflet, as indicated by molecular dynamics (MD) simulations, providing a pathway for membrane-embedded drugs. Hydrophobic substrates are expected to partition into the membrane and flip to the inner leaflet before entering the transporter through the TM5–TM10 cleft (**step II**). MD simulations indicate that substrate binding in the inward-open conformation is transient and unstable. Due to a limited energy barrier, a rigid-body transition to the outward-open state can occur (**step III**), enabling formation of a high-affinity binding site. For ethidium, this site involves residues W149, S153, M287, M380, E407 and D411, and requires local rearrangement of TM5. This stabilizes the outward-open conformation (**step IV**), in which the binding pocket is accessible to the extracellular medium. The extracellular environment is more acidic than the cytoplasm (schematized by blue circles representing protons), enabling protonation of the binding site, in particular residue E407, whose pKa is elevated in the outward-open state (**step V**). Protonation weakens substrate affinity, leading to its release to the extracellular side, thus achieving extrusion. Finally, consistent with an antiport mechanism, the proton is translocated across the transporter and released into the cytoplasm (**state VI**), thereby enabling return to the resting inward-open state; the molecular details of this step remain to be elucidated.

## Notes

https://github.com/OMaraLab/QacA_Structural_2026

### Competing Interest Statement

The authors have declared no competing interest.

### Summary of Updates

Corresponding authors added. Page numbers added.

## References

1. Murray, C.J. et al. (2022) Global burden of bacterial antimicrobial resistance in 2019: a systematic analysis. The lancet 399 (10325), 629–655.

2. Li, X.-Z. et al. (2015) The challenge of efflux-mediated antibiotic resistance in Gram-negative bacteria. Clinical microbiology reviews 28 (2), 337–418.

3. Brown, M.H. and Skurray, R.A. (2001) Staphylococcal multidrug efflux protein QacA. Journal of molecular microbiology and biotechnology 3 (2), 163–170.

4. Mancuso, G. et al. (2021) Bacterial antibiotic resistance: the most critical pathogens. Pathogens 10 (10), 1310.

5. Dashtbani-Roozbehani, A. and Brown, M.H. (2021) Efflux pump mediated antimicrobial resistance by staphylococci in health-related environments: challenges and the quest for inhibition. Antibiotics 10 (12), 1502.

6. Saier Jr, M.H., et al. (2021) The transporter classification database (TCDB): 2021 update. Nucleic acids research 49 (D1), D461–D467.

7. Drew, D. et al. (2021) Structures and general transport mechanisms by the major facilitator superfamily (MFS). Chemical reviews 121 (9), 5289–5335.

8. Martens, C. et al. (2016) Lipids modulate the conformational dynamics of a secondary multidrug transporter. Nature structural & molecular biology 23 (8), 744–751.

9. Debruycker, V. et al. (2020) An embedded lipid in the multidrug transporter LmrP suggests a mechanism for polyspecificity. Nature structural & molecular biology 27 (9), 829–835.

10. Wang, S. et al. (2024) Structures of the Mycobacterium tuberculosis efflux pump EfpA reveal the mechanisms of transport and inhibition. Nature Communications 15 (1), 7710.

11. Majumder, P. et al. (2023) Cryo-EM structure of antibacterial efflux transporter QacA from Staphylococcus aureus reveals a novel extracellular loop with allosteric role. The EMBO Journal 42 (16), e113418.

12. Quistgaard, E.M. et al. (2016) Understanding transport by the major facilitator superfamily (MFS): structures pave the way. Nature Reviews Molecular Cell Biology 17 (2), 123–132.

13. Heng, J. et al. (2015) Substrate-bound structure of the E. coli multidrug resistance transporter MdfA. Cell research 25 (9), 1060–1073.

14. Masureel, M. et al. (2014) Protonation drives the conformational switch in the multidrug transporter LmrP. Nature chemical biology 10 (2), 149–155.

15. Paulsen, I.T. et al. (1996) Proton-dependent multidrug efflux systems. Microbiological reviews 60 (4), 575–608.

16. Varela, M.F. et al. (2023) Functional Roles of the Conserved Amino Acid Sequence Motif C, the Antiporter Motif, in Membrane Transporters of the Major Facilitator Superfamily. Biology 12 (10), 1336.

17. Wu, J. et al. (2008) Functional analyses reveal an important role for tyrosine residues in the staphylococcal multidrug efflux protein QacA. BMC microbiology 8, 1–7.

18. Mitchell, B.A. et al. (1999) Bioenergetics of the staphylococcal multidrug export protein QacA: identification of distinct binding sites for monovalent and divalent cations. Journal of Biological Chemistry 274 (6), 3541–3548.

19. Dashtbani-Roozbehani, A. et al. (2023) The role of TMS 12 in the staphylococcal multidrug efflux protein QacA. Journal of Antimicrobial Chemotherapy 78 (6), 1522–1531.

20. Majumder, P. et al. (2019) Dissection of protonation sites for antibacterial recognition and transport in QacA, a multi-drug efflux transporter. Journal of molecular biology 431 (11), 2163–2179.

21. Hassan, K.A. et al. (2008) Analysis of tryptophan residues in the staphylococcal multidrug transporter QacA reveals long-distance functional associations of residues on opposite sides of the membrane. Journal of bacteriology 190 (7), 2441–2449.

22. Valley, C.C. et al. (2012) The methionine-aromatic motif plays a unique role in stabilizing protein structure. Journal of Biological Chemistry 287 (42), 34979–34991.

23. Fluman, N. et al. (2014) Export of a single drug molecule in two transport cycles by a multidrug efflux pump. Nature communications 5 (1), 4615.

24. Bas, D.C. et al. (2008) Very fast prediction and rationalization of pKa values for protein–ligand complexes. Proteins: Structure, Function, and Bioinformatics 73 (3), 765–783.

25. Senćanski, M. et al. (2015) Theoretical insight into sulfur–aromatic interactions with extension to D 2 receptor activation mechanism. Structural Chemistry 26, 1139–1149.

26. Xie, P. et al. (2025) A fiducial-assisted strategy compatible with resolving small MFS transporter structures in multiple conformations using cryo-EM. Nature communications 16 (1), 7.

27. Yu, E.W. et al. (2003) Structural basis of multiple drug-binding capacity of the AcrB multidrug efflux pump. Science 300 (5621), 976–980.

28. Wilhelm, J. and Pos, K.M. (2024) Molecular insights into the determinants of substrate specificity and efflux inhibition of the RND efflux pumps AcrB and AdeB. Microbiology 170 (2), 001438.

29. Morgan, C.E. et al. (2021) Cryoelectron microscopy structures of AdeB illuminate mechanisms of simultaneous binding and exporting of substrates. MBio 12 (1), 10.1128/mbio.03690-20.

30. Ginn, S.L. et al. (2000) The TetA(K) tetracycline/H(+) antiporter from Staphylococcus aureus: mutagenesis and functional analysis of motif C. J Bacteriol 182 (6), 1492–8.

31. Hassan, K.A. et al. (2006) Functional effects of intramembranous proline substitutions in the staphylococcal multidrug transporter QacA. FEMS microbiology letters 263 (1), 76–85.

32. Nagarathinam, K. et al. (2018) Outward open conformation of a Major Facilitator Superfamily multidrug/H+ antiporter provides insights into switching mechanism. Nature communications 9 (1), 4005.

33. Bolhuis, H. et al. (1996) Multidrug resistance in Lactococcus lactis: evidence for ATP-dependent drug extrusion from the inner leaflet of the cytoplasmic membrane. The EMBO Journal 15 (16), 4239–4245.

34. Hanahan, D. (1983) Studies on transformation of Escherichia coli with plasmids. Journal of molecular biology 166 (4), 557–580.

35. Hahn, A. et al. (2013) The outer membrane TolC-like channel HgdD is part of tripartite resistance-nodulation-cell division (RND) efflux systems conferring multiple-drug resistance in the cyanobacterium Anabaena sp. PCC7120. Journal of biological chemistry 288 (43), 31192-31205.

36. Xu, Z. et al. (2006) Role of transmembrane segment 10 in efflux mediated by the staphylococcal multidrug transport protein QacA. Journal of Biological Chemistry 281 (2), 792–799.

37. Emsley, P. and Cowtan, K. (2004) Coot: model-building tools for molecular graphics. Acta crystallographica section D: biological crystallography 60 (12), 2126–2132.

38. Smart, O.S. et al. (2012) Exploiting structure similarity in refinement: automated NCS and target-structure restraints in BUSTER. Acta Crystallographica Section D: Biological Crystallography 68 (4), 368–380.

39. Pardon, E. et al. (2014) A general protocol for the generation of Nanobodies for structural biology. Nat Protoc 9 (3), 674–93.

40. Eberhardt, J. et al. (2021) AutoDock Vina 1.2. 0: New docking methods, expanded force field, and python bindings. Journal of chemical information and modeling 61 (8), 3891-3898.

41. Trott, O. and Olson, A.J. (2010) AutoDock Vina: improving the speed and accuracy of docking with a new scoring function, efficient optimization, and multithreading. Journal of computational chemistry 31 (2), 455–461.

42. Li, H. et al. (2005) Very fast empirical prediction and rationalization of protein pKa values. Proteins: Structure, Function, and Bioinformatics 61 (4), 704–721.

43. Powers, N. and Jensen, J.H. (2006) Chemically accurate protein structures: Validation of protein NMR structures by comparison of measured and predicted p K a values. Journal of biomolecular NMR 35, 39–51.

44. Porter, M.A., et al. (2006) Hydrogen bonding is the prime determinant of carboxyl pKa values at the N-termini of α-helices. Proteins: Structure, Function, and Bioinformatics 63 (3), 621–635.

45. Davies, M.N. et al. (2006) Benchmarking pK a prediction. BMC biochemistry 7, 1–12.

46. Stroet, M. et al. (2018) Automated topology builder version 3.0: Prediction of solvation free enthalpies in water and hexane. Journal of chemical theory and computation 14 (11), 5834–5845.

47. Hermans, J. et al. (1984) A consistent empirical potential for water–protein interactions. Biopolymers: Original Research on Biomolecules 23 (8), 1513–1518.

48. Bekker, H. et al., Gromacs-a parallel computer for molecular-dynamics simulations, 4th international conference on computational physics (PC 92), World Scientific Publishing, 1993, pp. 252-256.

49. Berendsen, H.J. et al. (1995) GROMACS: A message-passing parallel molecular dynamics implementation. Computer physics communications 91 (1-3), 43–56.

50. Van Der Spoel, D., et al. (2005) GROMACS: fast, flexible, and free. Journal of computational chemistry 26 (16), 1701–1718.

51. Abraham, M.J. et al. (2015) GROMACS: High performance molecular simulations through multi-level parallelism from laptops to supercomputers. SoftwareX 1, 19–25.

52. Huang, W. et al. (2011) Validation of the GROMOS 54A7 force field with respect to β-peptide folding. Journal of chemical theory and computation 7 (5), 1237–1243.

53. Heinz, T.N. et al. (2001) Comparison of four methods to compute the dielectric permittivity of liquids from molecular dynamics simulations. The Journal of chemical physics 115 (3), 1125–1136.

54. Humphrey, W. et al. (1996) VMD: visual molecular dynamics. Journal of molecular graphics 14 (1), 33–38.

55. Kabsch, W. and Sander, C. (1983) Dictionary of protein secondary structure: pattern recognition of hydrogen-bonded and geometrical features. Biopolymers: Original Research on Biomolecules 22 (12), 2577–2637.

56. Lau, A.M. et al. (2021) Deuteros 2.0: peptide-level significance testing of data from hydrogen deuterium exchange mass spectrometry. Bioinformatics 37 (2), 270–272.

